# Crystal structures of a far-red photoreceptor in different light-absorbing states: insights into spectral tuning and light signaling

**DOI:** 10.64898/2026.07.21.739654

**Authors:** Linta M. Biju, Zhong Ren, Anastasia Kraskov, Sepalika Bandara, Elham Norouzi Sardareh, Chongyao Wei, Aditya G. Rao, Igor Schapiro, Peter Hildebrandt, Xiaojing Yang

## Abstract

Cyanobacteriochromes (CBCRs) are bilin-binding photoreceptors with remarkable spectral versatility. Using phycocyanobilin (PCB) as a chromophore, CBCRs regulate diverse light-dependent processes in cyanobacteria, ranging from photosynthesis to chromatic acclimation. Although extensive studies have uncovered multiple spectral tuning mechanisms in bilin-binding proteins, recent structural studies of far-red CBCRs suggest the existence of additional tuning strategies in both the *15Z* and *15E* states. Here we report crystal structures of the representative far-red CBCR Anacy_2551g3 in three distinct light-absorbing states, all of which adopt a compact *all-syn* PCB conformation. These structures demonstrate that *15Z/15E* photoisomerization in Anacy_2551g3 involves minimal chromophore rotation relative to the GAF domain, in stark contrast to other characterized bilin-based photoreceptors. To investigate the molecular basis of its far-red absorption, we examined the protonation and tautomeric states of PCB in the Pfr state using resonance Raman (RR) spectroscopy and quantum mechanics/molecular mechanics (QM/MM) calculations. Comparisons of experimental and calculated RR spectra support a bilin lactam as the predominant tautomeric form in the Pfr state. Integrating structural, spectroscopic, computational and mutational analyses, we propose that specific protein-chromophore interactions play critical roles in modulating chromophore conjugation beyond bilin coplanarity. Structural analyses further suggest a signaling model in which light regulation by Anacy_2551g3 is mediated through reversible switching between a high-affinity Pfr state and a low-affinity Po state that does not involve large chromophore motions. Together, these results provide new insights into how protein-chromophore coupling governs spectral tuning and light signaling in bilin-based photoreceptors.

**Significance statement:** Bilins are widespread biological pigments that mediate photoreception, light harvesting, and photosynthesis across diverse light environments. In a phenomenon known as spectral tuning, the optical properties of bilin-binding proteins are profoundly influenced by protein-chromophore interactions. Mechanistic understanding of spectral tuning and light signaling is important not only for advancing fundamental knowledge of light-sensitive proteins but also for developing new engineering strategies in synthetic biology and biotechnology. Recently discovered cyanobacteriochromes (CBCRs) exhibit remarkable spectral diversity and structural versatility, providing excellent model systems for dissecting the mechanisms of bilin-based photoreceptors. By integrating crystallography, spectroscopy and computational methods, this work examines three distinct light absorbing states of a representative far-red CBCR. Our findings reveal previously unrecognized mechanisms of spectral tuning and light signaling, highlighting the critical roles of protein–chromophore coupling and electrostatic interactions in regulating photoreceptor function.

## Introduction

Bilins are widespread organic pigments that serve as chromophores in proteins involved in photoreception, light harvesting, and photosynthesis (1–4). Also known as linear tetrapyrroles, they consist of four pyrrole rings (denoted A-D) connected by three methine bridges, with propionate groups extending from rings B and C. Depending on their chemical structure and extent of conjugation, bilin chromophores absorb light in a broad spectral range spanning the blue to red regions of the visible spectrum. When bound within proteins, these highly malleable pigments engage in extensive interactions with their protein environment, further extending spectral sensitivity across the ultraviolet and visible regions (5). In light harvesting systems, phycobiliproteins incorporate bilins of different chemical identities at strategically positioned sites within the phycobilisome, enabling efficient and directional energy transfer to the photosynthetic reaction center in cyanobacteria (1, 6). In light-sensing proteins, phytochrome photoreceptors exploit bilin photoisomerization to initiate signaling through protein structural changes that ultimately alter enzyme activities and protein-protein interactions (3, 7, 8).

In bilin-based photoreceptors, including canonical phytochromes and the more recently discovered cyanobacteriochromes (CBCRs), the chromophore is embedded in the GAF domain through a thioether linkage and typically adopts an extended *15,anti* conformation (8–10). Extensive structural, biochemical and spectroscopic studies have established that the primary photochemical event in these photoreceptors is a *15Z,anti/15E,anti* photo-isomerization, regardless of the chromophore type, domain architecture, or evolutionary origin of the photoreceptor (11–16). However, recent studies have revealed a subfamily of CBCRs that adopt a highly compact, *all-syn* bilin conformation(13, 17). Specifically, we previously demonstrated that the phycocyanobilin (PCB) in the far-red CBCR from *Anabaena cylindrica* (denoted Anacy_2551g3 or 2551g3) adopts an *all-Z,syn* conformation in the far-red-absorbing (Pfr) state (17). Likewise, the crystal structure of red/green CBCR RcaE revealed an *all-syn* PCB in the red-absorbing (Pr) state, corresponding to the *15E* configuration (13). 2551g3 belongs to a recently identified clade of photoreceptors known as far-red CBCRs (18). RcaE, on the other hand, was the first CBCR characterized to function as a photoreceptor kinase that regulates complementary chromatic acclimation in cyanobacteria (19). The discovery of these structurally unusual chromophores challenges the prevailing framework for bilin photochemistry and presents a unique opportunity to re-examine the molecular mechanisms underlying spectral tuning, photoconversion, and long-range signaling in bilin-based photoreceptors.

2551g3 is a small photosensory GAF domain comprising approximately 180 residues from a much larger multi-sensor photoreceptor kinase protein, Anacy_2551 (Fig. S1A). In solution, 2551g3 undergoes reversible photoconversion between the Pfr state and the orange-absorbing (Po) state (18) (Fig. S1B). In the Pfr structure, the *all-Z,syn* PCB chromophore engages several protein-chromophore interactions not commonly found in other phytochrome photoreceptors (17). Most notably, PCB adopts a highly twisted conformation distinct from the typical M- or P-configuration of free *all-syn* bilins in solution or protein-bound states (20, 21). The chromophore is sandwiched between two hallmark residues, Leu944 and Glu914, which are conserved among far-red CBCRs. The hydrophobic Leu944 on the α-face of PCB appears to force all four pyrrole rings towards the β-face, where they engage direct interactions with the acidic residue Glu914 (17). This unusual geometry challenges the widely accepted coplanarity model, which predicts that increased chromophore twisting reduces effective conjugation, and consequently shifts absorption toward shorter wavelengths (22). To explain the extreme far-red absorption of 2551g3, we previously proposed two distinct mechanisms. One involves lactam/lactim tautomerization of rings A or D, whereas another implicates electrostatic interactions between Glu914 and ring D (17). However, important aspects of these hypotheses remain unresolved. In particular, the protonation and tautomeric states of PCB in the Pfr and Po states have not been rigorously examined.

Acid denaturation experiments on 2551g3 and related systems establish that far-red CBCRs undergo photoconversion via *15Z/15E* photoisomerization (17, 18). Interestingly, members of the far-red CBCR family can populate two distinct *15E* states, denoted the *15E*-Po and *15E*-Pr states (17, 18). Several single mutants of 2551g3 display mixtures of these two states upon photoconversion, although their relative populations vary considerably among mutants (17). Extensive structural studies of canonical phytochromes and CBCRs support a consensus model in which the ring D flip is accompanied by a large-scale rotational motion of the entire chromophore relative to the protein scaffold (11, 12, 15, 16). This “flip-and-rotate” model also appear to apply to RcaE despite its *all-syn* PCB conformation. Comparison between the Pr and Pg structures of RcaE have revealed a substantial chromophore reorientation relative to the protein framework during photoconversion (13, 14). In the absence of structural information for the *15E* state, it is unclear how 2551g3 accommodates *15Z/15E* photoisomerization in a highly restrictive protein pocket enabling photoconversion with a large spectral shift. In particular, it remains an open question whether Pfr/Po photoconversion in 2551g3 follows the canonical flip-and-rotate mechanism or proceeds through a fundamentally different pathway.

In naturally occurring proteins, far-red CBCRs functions as photosensory modules that are connected to diverse effector domains within multi-domain photoreceptors, where light-induced structural changes alter enzymatic activities through allosteric mechanisms (8) (Fig. S1A). Far-red CBCRs are also attractive tools in optogenetic applications because their action spectra fall within the optical window (650-950 nm), which enables non-invasive manipulation in deep tissues (23). In addition, the compact size and modular architecture of CBCRs offers significant advantages for the development of synthetic biology tools through protein engineering (2, 24).

In this work, we present crystal structures of 2551g3 corresponding to three distinct light-absorbing states: *15Z*-Pfr, *15E*-Pr, and *15E*-Po. Comparative structural analyses reveal key protein-chromophore interaction networks associated with each state and highlight the hitherto unrecognized role of electrostatic interactions in spectral tuning. We further investigate the protonation and tautomeric states of the bilin chromophore in the Pfr and Po states using resonance Raman (RR) and infrared (IR) spectroscopy. Guided by QM/MM calculations and comparisons with representative CBCR systems, we re-examine several prevailing models of spectral tuning in bilin-binding proteins. With implications extending beyond far-red CBCRs, our findings challenge several widely accepted notions regarding spectral tuning and light signaling in bilin-based photoreceptors. More broadly, this work provides new insights into how protein–chromophore coupling governs both the optical properties of bilins and the mechanisms by which light signals are transmitted through GAF-mediated photoreceptors.

## Results

### 2551g3 dimer structure shows mixed *15Z*-Pfr and *15E*-Pr states

Cryo-crystallography datasets collected from the green-colored 2551g3 crystals using conventional methods did not yield diffraction resolution beyond 3.0 Å, despite harvesting and freezing under controlled light conditions. While the corresponding 2Fo-Fc maps show clear electron density for the protein moiety, the chromophore region is poorly resolved, and only the rings A-C are visible (Fig. S1). These observations suggest that 2551g3 crystals are highly sensitive to ambient light, and the light sensitivity results in a highly mobile ring D in the crystalline state even at cryogenic temperatures. To capture the “true” dark-adapted state, we obtained a new crystal form for 2551g3 and adopted a room-temperature serial crystallography approach. This platform enables automated serial *in situ* Laue data collection from thousands of crystals grown on chips (25, 26). Crystals were identified under safe infra-red illumination and diffracted at room temperature (Fig. S2A). The final dataset at 2.4 Å resolution was processed from 3936 individual Laue images using Precognition. In the P4_2_ space group, the structure contains two 2551g3 molecules per asymmetric unit, forming a “handshake” dimer similar to the previously reported Pfr structure (Fig. 1A).

**Figure 1.**
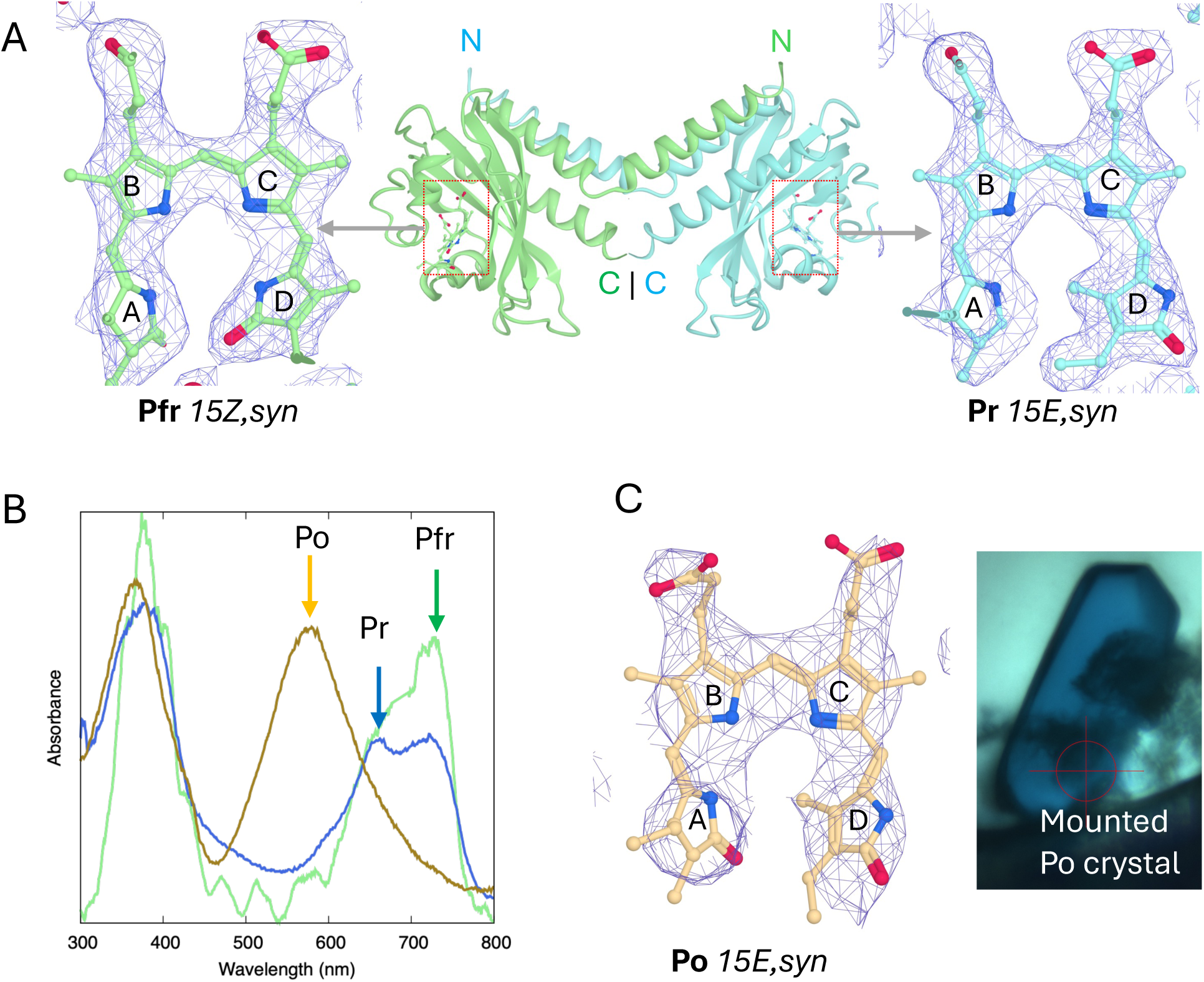
Crystal structures of 2551g3 in three light-absorbing states. **A)** Mixed Pfr/Pr states within the 2551g3 dimer. In the asymmetric unit, the PCB chromophore exhibits an *all-syn*,*15Z* conformation (green) in one protomer while adopting an *all,syn*,*15E* state (cyan) in the other. Electron densities of the PCB chromophore are shown in 2*F*o-*F*c map (contoured at 1.5σ). **B)** Single-crystal spectra of three different 2551g3 crystal forms: dark-adapted crystal in P4_2_22 space group (green, Bandara et al. 2021), dark-adapted crystal in P4_2_ space group (blue, this work) and light-adapted crystal in I222 space group (yellow, this work). **C) Left:** 2*F*o-*F*c map (contoured at 1.0σ) of the *15E*-Po state shows an *all-syn* PCB conformation. **Right:** A blue-colored Po crystal mounted on a cryo-loop used for cryo-crystallographic data collection.

Due to its high redundancy, the room-temperature Laue dataset yields electron density maps of exceptional clarity, enabling confident modeling of the entire chromophore (Fig. 1A). Both chromophores within the same dimer adopt an *all-syn* conformation. However, structure refinement reveals two distinct ring D configurations, with one protomer in the *15Z* state and the other in the *15E* state (Fig. 1A). In the *15Z* structure, all four pyrrole nitrogen atoms are within the hydrogen-bonding distance (2.6-3 Å) of the hallmark residue Glu914 (Fig. 2A). In the 1*5E* structure, ring D is flipped outwards, with its carbonyl group forming hydrogen bonds (3.1 Å) with the side chain of Glu946, a highly conserved residue at the C-terminus of a short helix at the α-face of PCB (Fig. 2A). In both states, the ring C propionate is coordinated by Tyr947 and His989 while the ring B propionate is tightly associated with Lys956, Arg930 and Tyr924 (Fig. 2A). Beyond the ring D flip, the two states differ markedly in ring D tilt relative to the remaining PCB moiety. In addition, His989 shows a nearly parallel rotamer relative to the ring C propionate in the *15Z* state while adopting a nearly vertical orientation in *15E* (Fig. 2A).

**Figure 2.**
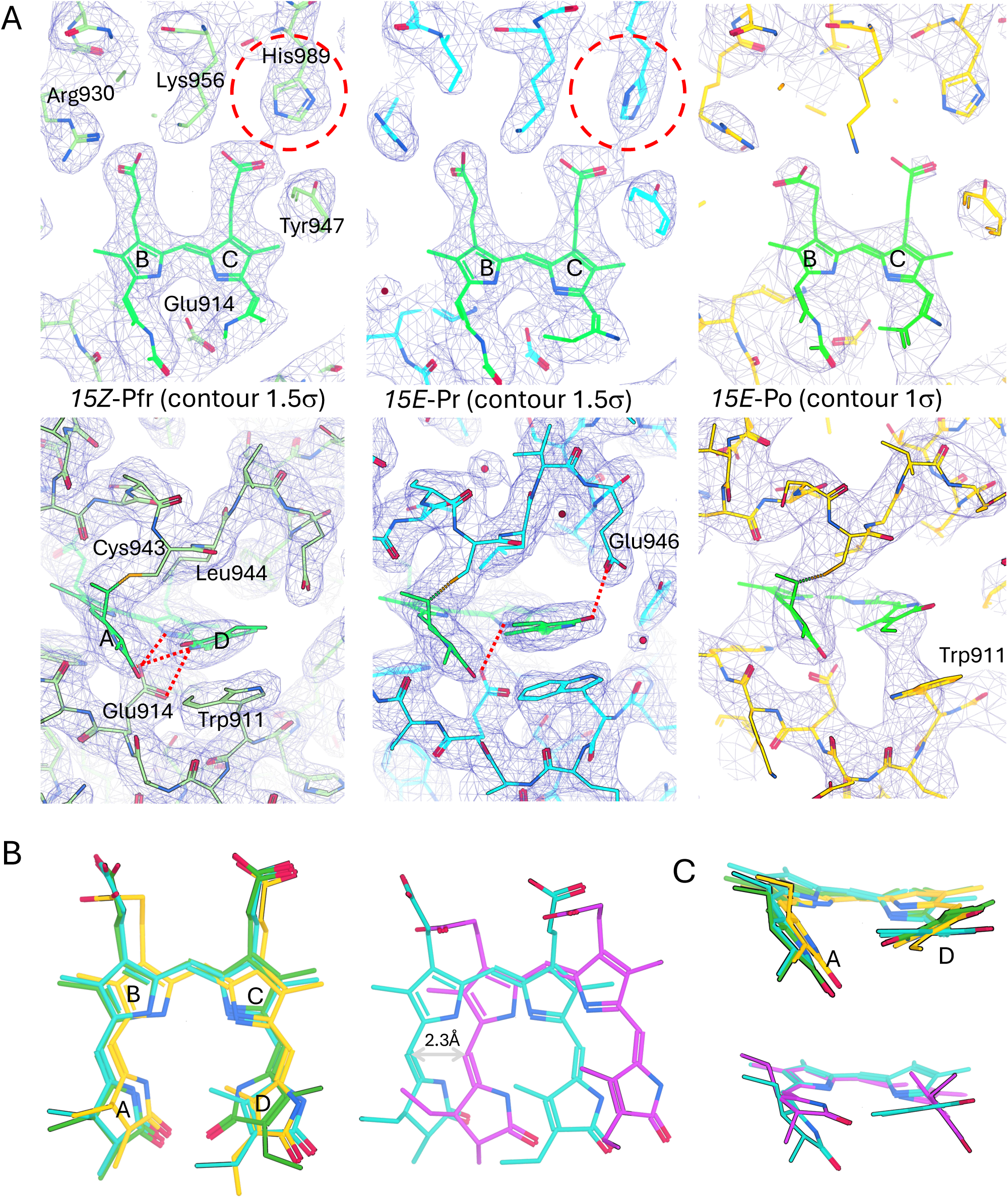
Comparisons of the chromophore conformation and protein environment between *15Z*-Pfr (green), *15E*-Pr (green) and *15E*-Po (yellow) states. **A)** Top: face-on views of the chromophore highlight the differences in protein-chromophore interactions between the propionate groups and charged residues (Arg930, Lys956 and His989) extending from the central β-strands of the GAF core domain. Bottom: edge-on views of ring D show the chromophore environment including the covalent anchor Cys943, two hallmark residues Leu944/Glu914 that sandwich the PCB, and Trp911 in stacking interactions with ring D. The *15Z*-Pfr state (left) features short hydrogen bonds (red dash lines) between the side chain of Glu914 and the pyrrole nitrogen of rings C and D. In the *15E*-Pr state (middle), the flipped ring D forms a hydrogen bond with Glu946 via its carbonyl group. In the *15E*-Po state (right), ring D is primarily stabilized by Trp911 evidenced by their connected densities. For clarity, the chromophores in all states are colored in green. **B)** Superposition of the chromophores based on the protein structure alignment (face-on view). **Left**: A slight chromophore shift (∼0.6 Å) relative to the protein frameworkbetween the Pfr and Po structures. **Right**: A significant chromophore shift (∼2.3 Å) between the *15E*-Pr structures of 2551g3 (cyan) and RcaE (magenta, PDB ID: 7CKV) (magenta). **C)** Edge-on views are shown to highlight the differences in ring twist and ring D tilt. **Bottom**: Superposition of 2551g3 (cyan) and RcaE (re-aligned according to ring B of the chromophore) shows that these two *15E*-Pr structures exhibit distinct ring D disposition.

To determine the light-absorbing state represented by this *15E* structure, we analyzed single-crystal absorption spectra collected from 2551g3 crystals (Fig. 1B). In contrast to the previously reported Pfr crystal, this new crystal form shows a prominent absorption band at ∼655-nm in addition to the 728-nm peak characteristic of the Pfr state. This Pr-like band closely matches the *15E*-Pr state previously identified in several far-red CBCRs that photoconvert to the *15E*-Pr instead of the *15E*-Po state (Fig. S3A) (17, 18). Upon photoconversion, several mutants of far-red CBCRs including 2551g3 and 4718g3 exhibit pronounced peaks around 650-690 nm, attributed to the formation of the 1*5E*-Pr state (Fig. S3A)(17). To confirm whether the *15E* state indeed exists in crystals, we carried out acid denaturation experiments on solution samples as well as dissolved crystals under different illumination conditions. The denatured spectra were subjected to a least-squares fitting procedure that performs both composition analysis and baseline corrections (see Methods). Not surprisingly, all tested samples show mixed *15Z*/*15E* states with different ratios depending on light treatment (Fig. S3B). While pre-illumination with 550-nm light resulted in a predominant *15Z* population, the dissolved and denatured 2551g3 sample collected from dark-adapted crystals gave rise to a nearly 1:1 composition (52% *15Z* vs. 48% *15E*), consistent with our crystallography observations. Based on structural data and comparative spectral analysis, we conclude that the new crystal form has captured a conformational state where the *15Z*-Pfr and *15E-Pr* states co-exist within the same dimer. Such heterodimeric assemblies with mixed *15Z/15E* configurations have been reported in other bilin-binding photoreceptor proteins, both in crystalline form and in solution (27, 28). It is noteworthy that this Pr band is not detected in static spectroscopic measurements of 2551g3 solution samples (Fig. S1B). However, a transient species associated with 645-nm absorption has been observed for 2551g3 on the 100-ps time scale in time-resolved spectroscopic studies of the *15E* state (29). It is plausible that the *15E*-Pr state observed in this work arises from specific protein-chromophore interactions fortuitously stabilized by crystal lattice in the P4_2_ space group.

### The Po structure reveals weakened association between PCB and the GAF core

2551g3 remains photoactive in the crystalline state. In a time series single-crystal spectroscopy experiment, the 588-nm absorption band associated with the Po state is neglectable in dark-adapted crystals but increases markedly upon red light illumination (Fig. S4A). Consistently, 2551g3 crystals undergo an apparent color change from green to blue (Fig. S4B). However, these illuminated crystals became jelly-like and failed to yield diffraction of sufficient quality, suggesting lattice disorder induced by light-driven structural changes. To obtain crystals suitable for structural characterization of the Po state, we pre-illuminated 2551g3 in solution prior to crystallization under continuous white light. This approach only produced a limited number of blue-colored crystals used for X-ray data collection (Fig. 1C). These Po crystals were difficult to reproduce and diffracted only to 3.5-4.0 Å resolution (Table S1). Nevertheless, they offered a rare opportunity to examine the Po state, as confirmed by single-crystal spectroscopy (Fig. 1B).

In the I222 space group, the Po structure contains five 2551g3 molecules in the asymmetric unit (Fig. S5). Three molecules (chains A-C) form crystallography dimers with their respective symmetry mates, while the remaining two (chains D and E) form a non-crystallography dimer. Chain E exhibits overall weak density, likely due to significantly increased molecular flexibility within the crystal lattice. Despite this unusual packing, all modes of dimerization are consistent with the “handshake” dimer observed in the dark-adapted structure (Fig. S5A). There are no major structural differences among five protomers in the GAF core domain. However, substantial variation is observed in the disposition of the GAF-α1 helix relative to the core domain (Fig. S5B). Although the resolution is limited, the electron density is of sufficient clarity to model an *all-syn* PCB conformation in chains A-D. Based on the single-crystal spectra showing the Po state, all five chromophores are assigned a *15E,syn* configuration. This Po structure was refined to 3.6 Å resolution with a final *R*-factor and *R*-factor of 0.246 and 0.297, respectively.

Compared to the *15Z*-Pfr and *15E*-Pr structures, coupling between PCB and the GAF core appears significantly weakened in the 1*5E*-Po structure. The ring-C propionate is primarily stabilized by Tyr947, while the ring B propionate forms a branched salt bridge with Arg930 and Asp938. Notably, no electron density above the 1σ level is observed for the side chain of Lys956 in any protomer, indicating loss of its interactions with both propionates (Fig. 2A). Consistent with this, electron density connectivity shows that His989 no longer engages in close contacts with the ring C propionate while Glu946 shifts away from ring D (Fig. 2B). Concomitantly, the density of ring D becomes diffused and merges towards the indole ring of Trp911 (Fig. 2A). Collectively, these features suggest increased conformational flexibility in the Po state, which likely underlies the difficulty in obtaining high-quality Po crystals.

The side chains of Arg930, Lys956 and His989 originate from the central β3, β4 and β5 strands of the GAF core domain, respectively (Fig. 2A). Substitution of these residues with Leu completely abolishes PCB binding, underscoring their critical role in chromophore association (17). In addition, pH titration studies of far-red CBCRs show that the Po state can be induced at alkaline pH conditions in the absence of light (17). Taken together, we postulate that the Po structure represents a conformational state where the GAF core loosens its interactions (i.e. affinity) with the chromophore. As a result, steric effects from surrounding bulky residues become dominant, leading to a highly twisted chromophore conformation that gives rise to the Po state (Fig. 2B). This interpretation is consistent with the published mutational data showing that substitution of Trp residues surrounding the chromophore selectively affect the Po state, while leaving the Pfr state largely unchanged (17).

### Photoconversion of 2551g3HK chimera mediates light-dependent kinase regulation

Acid denaturation experiments have established that Pfr/Po photoconversion in 2551g3 involves *15Z*/*15E* isomerization (Fig. S3B). This study shows that both the Pfr and Po structures adopt an *all,syn* PCB conformation, with the overall disposition of the chromophore relative to the GAF domain largely unchanged except the flip of ring D. In cryo-crystallography studies where it is difficult to protect light-sensitive crystals from light exposure, we observed well defined electron density for rings A-C while ring D is highly disordered, further supporting that ring D motion is the primary source of light sensitivity in 2551g3 (Fig. S1C). This finding, however, contrasts with the prevailing “flip-and-rotate” model proposed for phytochrome-family photoreceptors (11, 16), raising the question of whether 2551g3 can mediate light signaling without substantial bilin rotation. To address this, we constructed a chimeric protein where the photosensory domain of 2551g3 is fused to the effector domains of PPHK, using full-length Anacy2551 as a template (Fig. 3A). Using the C-terminal receiver domain (c-REC) of PPHK as a readout, we perform histidine kinase assays under different light conditions. The resulting 2551g3HK chimera exhibits clear light-dependent regulation of kinase activity, with the Po state showing higher kinase activity than the Pfr state (Fig. 3B).

**Figure 3.**
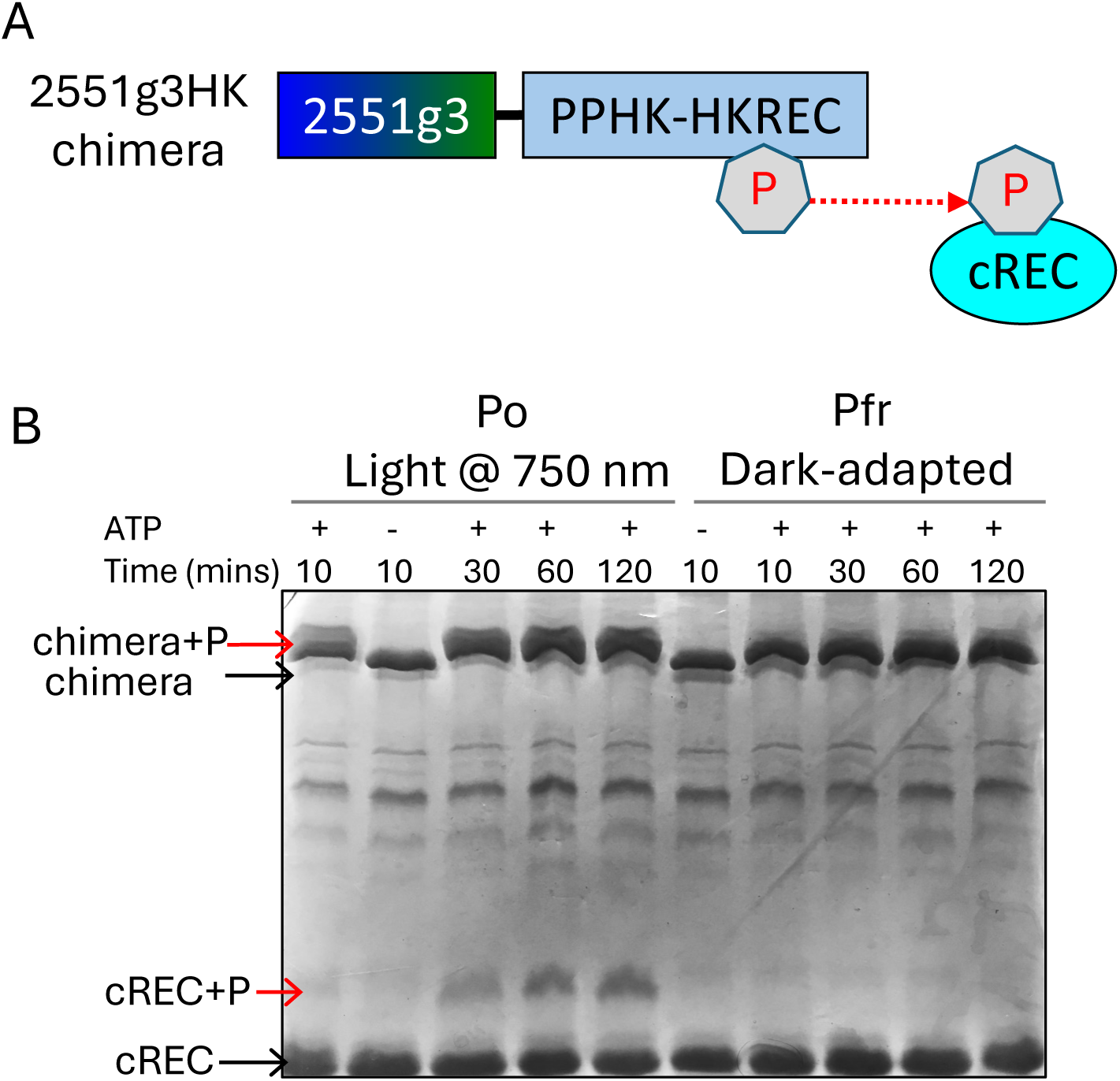
Engineering of 2551g3HK chimera and light regulation of histidine kinase. **A)** Domain architecture of 2551g3HK chimera where 2551g3 is fused to the effector domains (HK: histidine kinase in blue; cREC: C-terminal receiver domain in cyan) of a dual sensor histidine kinase PPHK (Shin et al. PNAS, 2019). cREC is the corresponding reporter protein that receives a phosphoryl group (letter P in hexagon) from phosphorylated HK. **B)** Light-dependent regulation of 2551g3HK chimera. In a PhosTag-based gel-shift assay, phosphorylated bands (red arrows) migrate slower than unphosphorylated bands (black arrows). Reactions quenched after 10, 30, 60, 120 minutes incubation after adding ATP are probed for the HK autophosphorylation (upper bands) and cREC phosphorylation (lower bands). The 2551g3HK chimera exhibits upregulated kinase activity in the Po-enriched state than the dark-adapted Pfr-enriched state.

It remains unclear how the Pfr/Po photoconversion alters the protein moiety to confer allosteric regulation of the C-terminal kinase domain. While addressing this mechanism will require further structural studies of full-length proteins or different constructs, the present work reveals a subtle yet remarkable difference between the Pfr and Po structures in terms of chromophore-protein interactions. In the Pfr state, the PCB chromophore maintains a tight association with the GAF core, whereas in the Po state, it appears partially disengaged, accompanied by disruption of key salt bridges around the propionates, despite remaining covalently linked in both states. Beyond the chromophore pocket, notable structural differences between the *15Z*-Pfr and *15E*-Pr states reside in a large β4-β5 loop insertion spanning residue 963-980 as revealed by difference distance matrix between the main chain atoms of these two structures (Fig. S6A). Characteristic to far-red CBCRs, this loop is part of the tri-loop junction, a unique structural feature that bridges the chromophore to the GAF-α3 helix at the dimer interface (17). Structural comparison between *15Z*-Pfr and *15E*-Po structures reveal more extensive changes associated with the tri-loop junction in addition to the varying disposition of GAF-α1 helix (Fig. S6B). It is plausible that subtle changes in the chromophore are transmitted via the tri-loop junction to the signaling helices at the dimer interface, thereby altering the remote HK activity via the shared central helical spine.

### Spectral tuning in the *15E,syn* state

We compared the Po and Pr structures of 2551g3 with the Pr structure of RcaE (PDB ID: 7ckv) (Fig. 2B). All three structures feature a *15E,syn* PCB sandwiched between two hallmark residues (i.e. α-facial Leu and β-facial Glu). However, their chromophores differ in their ring twist and ring D disposition. Compared to *15E*-Pr of 2551g3, the PCB in the *15E*-Po state exhibits increased twisting in the C-D methine bridge, with the ring D carbonyl tilted towards the α face (Fig. 2B). In contrast, the RcaE-Pr structure displays a PCB conformation with nearly co-planar rings A-C while its ring D carbonyl points towards the β face (13, 17). While the ring twist model may explain the 60-nm absorption shift between the Po and Pr states of 2551g3, it does not fully account for the differences between the Pr structures of 2551g3 and RcaE (Fig. 2B, Fig. 1B), suggesting that additional tuning mechanisms are involved.

In the RcaE-Pr structure, ring D is located in a largely hydrophobic pocket (13, 14). In contrast, the ring D carbonyl directly interacts with Glu946 in the 2551g3-Pr structure (Fig. 2A). Glu946 is conserved among far-red CBCRs in the far-red/X cluster, where X represents different *15E* states (17). This interaction is only observed in the new crystal form of 2551g3 (P4_2_), which exhibits mixed Pfr/Pr states according to single-crystal spectroscopy (Fig. 1B). In contrast, in the previous crystal form (P4_2_22) showing a nearly 100% pure Pfr state, the side chain of Glu946 appears highly disordered. To investigate the role of Glu946, we substituted this residue with Leu in wildtype 2551g3 and the W940L mutant that features a prominent *15E-*Pr species. While the single mutant E946L does not affect the Pfr/Po photoconversion, the double mutant E946L/W940L alters the Pr/Po distribution in the *15E* state compared to the single mutant W940L (Fig. S3A). These results suggest that the *15E*-Pr structure is not an authentic intermediate in the Pfr/Po photoconversion pathway. In other words, it may represent a fortuitous photoproduct arising from changes in the local environment of ring D, due to either mutations or crystal packing effects (17). This structure along with single crystal spectra, nevertheless, allows us to examine the tuning effects exerted by specific interactions with ring D. On one hand, Glu946 stabilizes ring D via hydrogen bonding, thereby limiting extensive ring twist across the C-D methine bridge. On the other hand, its interaction with the ring D carbonyl may exert an inductive effect that counteracts the blue shift associated with the resonance changes induced by ring twisting. Notably, similar interactions between the ring D carbonyl and a polar/acidic residue are commonly observed in phytochrome Pr structures (9, 30). It is important to note that 2551g3 is not an ideal system to test the role of Glu946 because the *15E*-Pr state is not detectable in solution using standard methods.

In the *15E* state, additional factors surrounding the chromophore also influence its absorption properties. Tyr947, which forms a short hydrogen bond with the ring C propionate, plays an important role in stabilizing the *15E*-Po state. Substitution of Tyr947 with Leu or Phe results in a pronounced shift from the *15E*-Po to the *15E*-Pr population (see Fig. S9 in Ref. (17)). Comparison between the Po and Pr structures suggests that a balance between Tyr947 and His989 directly affects the PCB association with the GAF core, which influences the extent of ring twist across the C-D methine bridge. The formation of *15E*-Po state is largely driven by steric effects imposed by Trp911 and other bulky residues in the chromophore pocket. Consistently, single mutations of aromatic residues, such as W911F, W925L, and W940L, selectively perturb the *15E*-Po state while leaving the *15Z*-Pfr state largely unaffected (17). Notably, the *15E*-Pr band becomes a major species in the W1350A and Y1386F mutants of 4718g3 (corresponding to Trp911 and Tyr947 of 2551g3, respectively) (17). Taken together, these results suggest that steric interactions play a dominant role in promoting the *15E*-Po state, in which the PCB chromophore is loosely coupled to the core β-sheet of the GAF domain (Fig. 2A).

### Resonance Raman spectroscopy probes chromophore structures in the Pfr and Po states

To investigate the chromophore configuration and protonation states of 2551g3 in solution, we measured the RR spectra of the Pfr and Po states. In the dark-adapted Pfr state, a prominent band at 1616 cm^−1^ is observed, which shifts to 1642 cm^−1^ upon photoconversion to the Po state (Fig. S7, S8). This mode is assigned to the C=C stretching of the C-D methine bridge based on comparisons with other bilin photoreceptors (31). The observed 26-cm^−1^ upshift of the C-D stretching mode from the Pfr to the Po state is consistent with photoinduced double bond isomerization between rings C and D.

The position of this mode between 1600 and 1640 cm^−1^ is characteristic of cationic bilins. In contrast, neutral tetrapyrroles exhibit marked different band patterns in this region, typically accompanied by weaker H/D isotope effects (32). In protonated bilins, the C-D methine bridge stretching mode includes minor contributions from N-H in-plane (NH ip) coordinates of adjacent pyrrole rings, resulting in small downshifts upon H/D exchange (Fig. S7). Consistently, 2551g3 shows pronounced isotope effects across the 1100-1600 cm^−1^ spectral region. In particular, the C-D stretching modes downshift by 7 and 12 cm^−1^ in the Pfr and Po states, respectively. These substantial isotopic effects provide strong evidence that the bilin chromophore of 2551g3 remains in a cationic form in both the Pfr and Po states.

We also collected the IR spectra in both states. The IR difference spectrum (Po-minus-Pfr) (Fig. S9) reveals pronounced features in the region in the 1680-1750 cm^−1^ region, corresponding to C=O stretching modes of the ring A and ring D carbonyl groups. The band pair at 1696/1710 cm^−1^ (Po/Pfr) is assigned to the ring D, whereas the pair at 1746/1726 cm^−1^ corresponds to ring A. The lower frequency of the ring D carbonyl mode reflects stronger coupling to the conjugated π-electron system, whereas the higher frequency of the ring A carbonyl suggests a more hydrophobic surrounding (33–36). Notably, the ring A carbonyl band shifts to 1735 and 1717 cm^−1^ in D_2_O for the Po and Pfr states, respectively. This relatively large deuteration-induced frequency downshift (9-11 cm^−1^) excludes assignment to a carboxyl side chain in either the PCB or the protein. Instead, this shift may result from coupling of NH ip coordinate to the ring A C=O stretching, indicating that the pyrrole nitrogen of ring A is protonated (36). The presence of two C=O stretching modes in both Pfr and Po states suggests that the bilin chromophore adopts a lactam isomer in both states, with a possible minor contribution from a ring A lactim form.

The IR difference signals in the 1610-1680 cm^−1^ region, corresponding to amide I modes, are not significantly stronger than those associated with chromophore carbonyl stretching (Fig. S9). This suggests that Pfr/Po photoconversion of 2551g3 does not involve major rearrangements in the protein moiety, consistent with our structural observation that the chromophore retains a *15-syn* configuration in both states. This finding contrasts with canonical phytochromes and CBCRs, where the bilin chromophore undergoes concerted “flip-and-rotate” motions relative to the protein framework during *15Z*/*15E* photoconversion (Fig. S10). For example, in the red/green CBCR RcaE, the Pr state adopts a compact *all,syn* conformation similar to that of 2551g3, whereas the Pg state exhibits an extended *15-anti* chromophore that is rotated by nearly 90 degree relative to the Pr state (13, 14, 37, 38)(Fig. S10B).

Despite the unique chromophore geometry of 2551g3, the RR spectra of the Pfr and Po states display vibrational patterns similar to those of canonical phytochromes. The hydrogen-out-of-plane (HOOP) modes occur at 813 cm^−1^ in Pfr and 812 cm^−1^ in Po, although their intensities differ from those observed in systems with *15-anti* chromophores (Fig. S8). This reflects a distinct distortion of the C-D methine bridge associated with the *all-syn* bilin configuration (31, 39). The primary difference lies in the NH ip modes of the pyrrole rings B and C. In canonical phytochromes, these modes are highly localized, typically giving rise to one RR-active and one IR-active band, with the RR-active feature appearing at 1550-1580 cm^−1^ in H_2_O and 1050-1070 cm^−1^ in D_2_O. In contrast, the NH ip coordinates of 2551g3 are distributed over multiple vibrational modes in both Pfr and Po states. In the Pfr state, three closely spaced modes are observed between 1508 and 1536 cm^−1^ in H_2_O (Fig. S7, S8). Upon H/D exchange, several modes, including a band at 1163 cm^−1^ exhibit strong ND ip character. For the Po state, assignments of distinct NH ip or ND ip modes are more challenging, due to even greater distributive character. This observation is consistent with the crystallographic structures. In the Pfr structure, rings B and C of the *all-syn* chromophore are not co-planar, and all four pyrrole N-H groups form strong hydrogen bonds with the acidic side chain of Glu914 (Fig. 2A, Fig. S13). In the *15E* state, rings A-C retain hydrogen bonding to Glu914 while ring D is further twisted with its lactam group “jammed” in a tight protein pocket (Fig. 2A, Fig. S13). By contrast, in canonical phytochromes, the pyrrole N-H groups of rings A-C are coordinated via the main-chain carbonyl group of the conserved Asp residue in the DIP motif while ring D in the *15-anti,E* chromophore engages additional hydrogen bonds with the Asp side chain (10).

### QM/MM calculations support the lactam tautomer in the Pfr state

To identify the predominant tautomeric form of the chromophore in the Pfr state, we performed QM/MM calculations based on the crystal structure of the *15Z* configuration. Raman spectra were calculated in both H_2_O and D_2_O for four possible cationic tautomers, differing in the enol/keto equilibria of rings A and D (Fig. 4). The calculated spectra show substantial differences in the 1580-1680 cm^−1^ region, which includes diagnostic C=C stretching modes of the methine bridges and pyrrole rings (Fig. 4A). All three lactim tautomers exhibit rather complex, multibanded features in this region, in contrast to the experimental spectrum, which contains a single, slightly asymmetric peak (Fig. 4B). By comparison, the calculated spectrum of the lactam closely reproduces the experimental data with a deviation of only 6 cm^−1^. This agreement extends to the H/D isotope effects, which are well captured by the lactam model but not by the lactim forms. Nevertheless, minor contributions from lactim-1 (up to 30%) or lactim-2 (up to 15%) cannot be excluded, as these mixtures remain compatible with the experimental RR spectra (Fig. 4D). Taken together, these results suggest that the bilin chromophore predominantly adopts a lactam tautomeric form in the Pfr state. This conclusion is further supported by the observed H/D isotope effects involving NH_ip modes between 1420 and 1560 cm^−1^, as well as the IR difference spectra, which show two C=O stretching bands characteristic of a lactam chromophore (Fig. S9).

**Figure 4.**
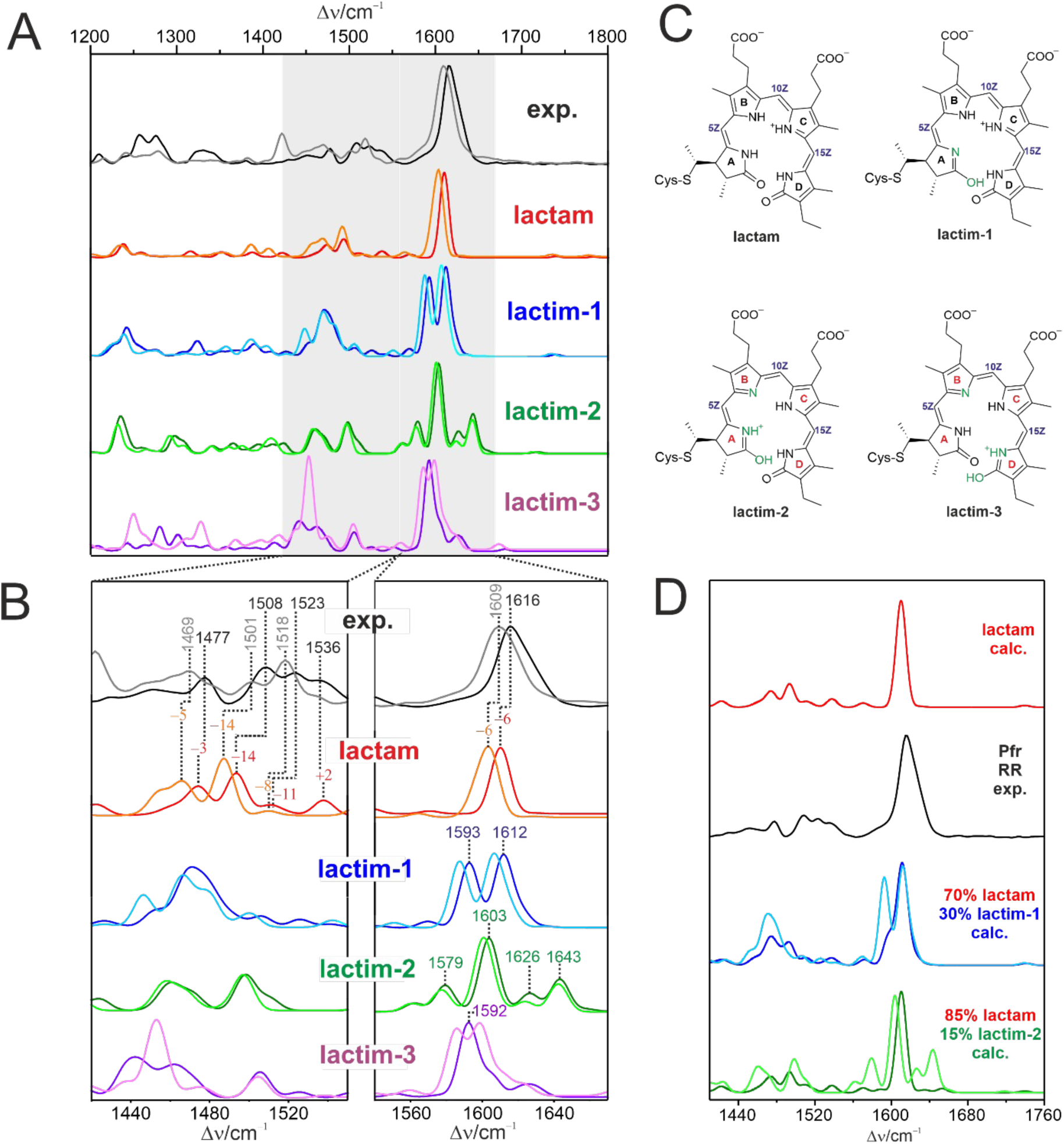
Experimental RR spectra and calculated Raman spectra in the Pfr state. **A)** Experimental RR spectra of Pfr in H_2_O (black) and D_2_O (grey) compared with the calculated Raman spectra of a lactam (red), lactim-1 (blue), lactim-2 (green), lactim-3 (magenta) in H_2_O and the same tautomers in D_2_O represented by the related pale colors. All calculated spectra were obtained by QM/MM and refer to the *ZZZsss* configuration. **B)** Expanded view of the experimental and calculated spectra from panel A in two diagnostic spectral regions. The intensity scale of the left panel is multiplied by a factor of 2.5. **C)** Possible tautomeric structures of the chromophore in the Pfr state. **D)** Experimental RR spectrum of Pfr (black) compared with the calculated Raman spectra of lactam (blue) and mixtures of the calculated spectra of lactam with 30% lactim-1 (magenta) and 15% lactim-2 (green). The pure calculated spectra of lactim-1 and lactim-2 are given in pale magenta and pale green, respectively.

To examine the chemical basis for pH-dependent behavior of 2551g3, we measured the RR spectra of both the Pfr and Po states over a pH range from 8.0 to 11.0 (Fig. S11). Although the overall spectral changes are modest, difference analysis between spectra collected at 8.0 and 11.0 allow decomposition into two components, which can be assigned to neutral and alkaline forms. The neutral component closely resembles the spectrum at pH 8, while the alkaline form contributes to approximately 45 - 50% at pH 11. The alkaline species closely matches the spectral features of deprotonated PCB (Fig. S12). Based on the relative contributions of these components, we estimate a pK_a_ of approximately 11 for chromophore deprotonation in both Pfr and Po states. This assignment is consistent with previously reported UV-vis absorption spectral data.

The corresponding IR difference spectra display two C=O stretching modes for both Pfr and Po across this pH range (Fig. S11), indicating that any lactim contribution remains minor even under alkaline conditions. Furthermore, no significant changes are observed in the amide I region at pH 10 – 11 compared to pH 8.0, suggesting that the overall protein structure remains largely unaffected by increasing pH.

## Discussion

### Light-sensitive crystals and in situ serial Laue crystallography at room temperature

Given the extreme light sensitivity of 2551g3 crystals, conventional cryo-crystallography methods did not yield electron density maps of sufficient quality for reliable structure interpretation. The present work was made possible by our newly developed serial diffraction platform that enables automated *in situ* Laue data collection at room temperature (25, 26, 40). In this approach, dark-adapted 2551g3 crystals are grown in sealed crystallization chips, visualized under infrared illumination, and diffracted directly at their growth locations (Fig. S2B). Because no crystal manipulation or cryo-cooling is required, this workflow minimizes unintended light exposure prior to data collection, and is therefore particularly well suited for highly photosensitive crystals (25, 26).

To mitigate X-ray radiation damage at room temperature, diffraction data were collected serially such that each crystal contributed only one or two diffraction images before being destroyed (Fig. S2C). This is evidenced by excellent densities observed for the covalent bond between Cys943 and ring A (Fig. 2A), which is prone to X-ray radiation damage in cryo-crystallography. Compared to conventional methods, serial crystallography requires large-scale scaling and merging of integrated diffraction intensities obtained from thousands of individual crystals (Fig. S2D). Consequently, merging and refinement statistics are often less favorable than those typically reported for single-crystal cryo-crystallography datasets (Table S1) (26, 40). As such, serial crystallography often exploits statistical metrics such as CC-halfs instead of traditional *R-*factors in reporting data quality. Compared to cryo-crystallography, room-temperature datasets are inherently associated with increased atomic mobility, which also contributes to higher R-factors. Regardless data statistics, our room-temperature serial Laue data produced electron density maps of exceptional quality, enabling confident structure interpretation of 2551g3 in a new crystal form (Fig. 1A). Similar to single particle cryoEM, the quality of our serial crystallography maps is largely attributable to high redundancy in merged datasets. With an average redundancy of approximately 420 in our final Laue dataset, the electron density maps generate remarkable clarity that led to the unexpected finding showing coexistence of *15E* and *15Z* chromophores within the same 2551g3 dimer, which is further supported by spectral composition analysis of denatured crystal samples (Fig. S3B). As the primary focus of this study is the structural biology of far-red CBCRs, the methodology advances underlying our serial Laue crystallography method will be described in detail elsewhere.

### Resonance and inductive effects in bilin spectral tuning

In a phenomenon known as spectral tuning, the absorption properties of a flexible organic pigment bound within a protein are determined not only by its intrinsic chemical structure but also by its surrounding protein environment (8, 41, 42). While the geometry of the chromophore is largely dictated by the binding pocket, its electronic properties can be modulated by both non-specific and specific protein-chromophore interactions through resonance and inductive effects. These mechanisms underlie remarkable spectral diversity of cyanobacteriochromes (CBCRs), which employ a common PCB chromophore within a conserved protein fold (2, 8, 43). However, the disposition of PCB relative to the protein moiety varies widely among CBCRs, although ring D typically adopts a 15,*anti* configuration (Fig. S9) (8, 13, 17). Previous work has demonstrated a strong correlation between chromophore coplanarity and absorption wavelength in both phycobiliproteins and CBCRs (17, 22). It supports the view that increased planarity enhances π-electron delocalization across the tetrapyrrole backbone, resulting in red-shifted absorption. Multiple spectral tuning strategies have been identified for PCB-binding proteins (Fig. S13). For instance, the two-Cys photocycle of TePixJ involves a photo-labile second covalent bond at C10, which introduces a chiral center and disrupts coplanarity between rings B and C (44, 45). In red/green CBCRs, blue-shifted absorption associated with Pr◊Pg photoconversion has been attributed to a “trapped twist” mechanism, in which bulky residues enforce torsion at the C-D methine bridge (46). Conversely, enhanced coplanarity of rings B-D has been proposed to explain the red-shift observed in the phycobiliprotein ApcD (22). Some CBCRs also exploit the PCB◊PVB isomerization, leading to blue-shifted absorption by disrupting the conjugation between rings A and B (8, 47). Collectively, these examples highlight the role of resonance effects in spectral tuning through direct modulation of the chromophore’s conjugation system (Fig. S13).

However, chromophore planarity alone cannot account for the extreme red-shift observed in 2551g3. To address this, we previously proposed two complementary mechanisms for the Pfr state (17). The first involves lactam/lactim tautomerization of PCB, motivated by the close proximity of the lactam groups of rings A and D in the highly twisted *all-syn* configuration. Computational studies by Noji et al. employing the QM/MM calculations with a polarizable continuum model (PCM) showed that the fully lactam form of PCB best reproduces the experimental absorption spectra of 2551g3 (48). This present work provides independent support for this conclusion. Our RR data, together with H/D isotope effects, indicate that the lactam tautomer most accurately accounts for the experimental spectra, whereas the lactim forms do not (Fig. 4). These findings establish the bilin lactam as the predominant tautomeric form in the Pfr state.

Noji et al. further proposed that far-red absorption of 2551g3 arises from increased orbital overlap between rings A and D PCB due to their close spatial proximity (48). Their QM/MM/PCM analysis also examined electronic contributions from residues surrounding the chromophore, revealing that polar/charged side chains exert modest but measurable tuning effects. In particular, residues interacting with propionate groups (Arg930, Tyr947, and Lys956) individually induce red-shifts of ∼4-10 nm, while the hallmark Glu914 contributes a red-shift by ∼4 nm (48). Interestingly, when similar strategies are applied to other CBCRs, a consistent trend has emerged: positively charged residues stabilizing the propionates tend to promote red-shift absorption, while acidic residues (Asp or Glu) interacting with pyrrole nitrogens at the β-face generally induce blue-shifts. The 2551g3-Pfr structure represents a notable exception as Glu914 engages in direct interactions with multiple pyrrole nitrogens, including that of ring D.

This observation reinforces the second tuning mechanism we previously proposed for far-red absorption in bilin-binding proteins (17). Specifically, we suggest that interactions between the ring D lactam and a strategically positioned acidic residue (Asp/Glu) plays a key role in extending absorption to far-red wavelengths, regardless of ring D orientation. While this mechanism has not been systematically evaluated across diverse systems by computational methods, it is supported by structural and spectral data obtained from 2551g3-Pfr (PCB), many phytochrome proteins (BV, PCB, or PΦB), and PcyA-I86D in complex with BV (17). In these structures, the carboxyl side chain of Asp or Glu acts as an electron-withdrawing group positioned near the ring D pyrrole nitrogen. We propose that this interaction, mediated through a short hydrogen bond, enhances electron delocalization across the tetrapyrrole system, effectively extending the conjugation length. Notably, this red-shifting effect contrasts with the blue-shifts predicted for interactions involving Glu914 and central pyrrole rings, which appear to reduce effective conjugation, highlighting the position-dependent nature of electrostatic tuning in bilin chromophores (48).

The unexpected *15E*-Pr structure captured in the new crystal form also features direct interaction between ring D and Glu946, analogous to the interaction between ring D and Glu914 in the *15Z*-Pfr structure. A key distinction, however, lies in the nature of these interactions. In the Pfr state, Glu914 forms a hydrogen bond with the pyrrole nitrogen of ring D, whereas in the Pr state, Glu946 interacts with the ring D carbonyl group. Because these interactions involve distinct electronic groups - electron-donating pyrrole nitrogen versus electron-withdrawing carbonyl, they are expected to exert different inductive effects, which may contribute to the substantial spectral difference between the Pfr and Pr states (728 nm vs. 655 nm). Notably, the overall degree of chromophore twisting is comparable between the *15Z*-Pfr and *15E*-Pr states, suggesting that PCB coplanarity is unlikely to be the dominant determinant of their spectral differences. We also attribute the spectral difference between the two *15E* states (655 nm vs. 588 nm) to a combination of resonance and inductive effects. Compared to the *15E*-Pr structure, the *15E*-Po structure exhibits a more pronounced chromophore twist and substantially weakened electrostatic interactions involving both the propionates and ring D. Collectively, these changes are expected to contribute to the blue-shifted absorption of the Po state. Clearly, further computational studies incorporating all three experimentally observed structures of 2551g3 would help delineate the respective contributions of resonance and inductive effects to spectral tuning.

Glu946 is highly conserved among far-red CBCRs. QM/MM calculations by Noji et al. suggest that Glu946 exerts a modest blue-shifting effect on the Pfr state, potentially through modulation of the polarity within the bilin-binding pocket (47). However, this prediction is not supported by our mutational analysis, as the E946L mutation does not significantly affect the Pfr state (Fig. S3A). Instead, substitution of Glu946 shifts the equilibrium between the two *15E* states. Specifically, the E946L mutation resulted in a marked shift from the *15E*-Pr state to the *15E*-Po state in the W940L/E946L double mutant.

### Photoconversion and light signaling mediated 2551g3

As a member in the phytochrome family, 2551g3 undergoes characteristic *15Z*/*15E* photo-isomerization. Indeed, our results suggest that light sensitivity of 2551g3 primarily involves ring D flip. However, the overall PCB disposition within the protein pocket remains unchanged. This is consistent with studies by IR difference spectroscopy, which did not detect substantial protein structural changes associated with photoconversion (Fig. S7). These observations contrasts sharply with the canonical “flip-and-rotate” model of phytochrome photoconversion, in which the bilin chromophore undergoes large-scale reorientation relative to the protein framework in addition to the ring D flip (Fig. S9) (11, 16, 49–52).

Despite the absence of pronounced chromophore rotation, Pfr/Po photoconversion clearly perturbs the protein structure of 2551g3. This is evidenced by the extraordinary light sensitivity of 2551g3 crystals. Crystals that were not adequately protected from light often diffracted poorly and exhibited highly disordered electron density around ring D (Fig. S1C). Moreover, illuminated crystals acquired a jelly-like texture, suggesting significant light-induced lattice disorder (Fig. S4B). Functional evidence for long-range signaling is provided by the light regulated histidine kinase activity of a chimeric protein in which the 2551g3 photosensory domain is fused to the kinase module of a different sensory protein (Fig. 3).

Consistently, considerable variation is observed for the GAF-α1 helix in terms of its orientation relative to the GAF core among the five crystallographically independent molecules in the Po structure (Fig. S5B). Tethered helices are commonly observed between juxtaposed GAF domains in phytochromes (Fig. 5A). Both crystallographic and cryoEM studies have revealed substantial light-induced rearrangements involving the GAF-α1 and GAF-α3 helices at the dimer interface (10, 27, 28, 49, 53). This recurring structural motif is thought to play a central role in mediating long-range signaling in modular kinase receptors. In this work, structural comparisons based on difference distance matrices reveal that largest structural differences between different light-absorbing states of 2551g3 are associated with the tri-loop junction, a structural feature unique to far-red CBCRs, which bridges the chromophore pocket to the central helical spine at the dimer interface (17) (Fig. S6, Fig. 5). We therefore propose that a similar signaling mechanism accounts for light regulation by 2551g3. Specifically, the tri-loop junction may offer a plausible pathway that facilitates signal transmission from the chromophore to GAFα3, a long signaling helix shared with the output histidine kinase (HK) domain (Fig. 5A). Clearly, further studies on larger constructs that contains both sensor and effector domains are needed to uncover the structural and mechanistic details to explain how far-red CBCRs achieves long-range signaling without the significant chromophore motions.

**Figure 5.**
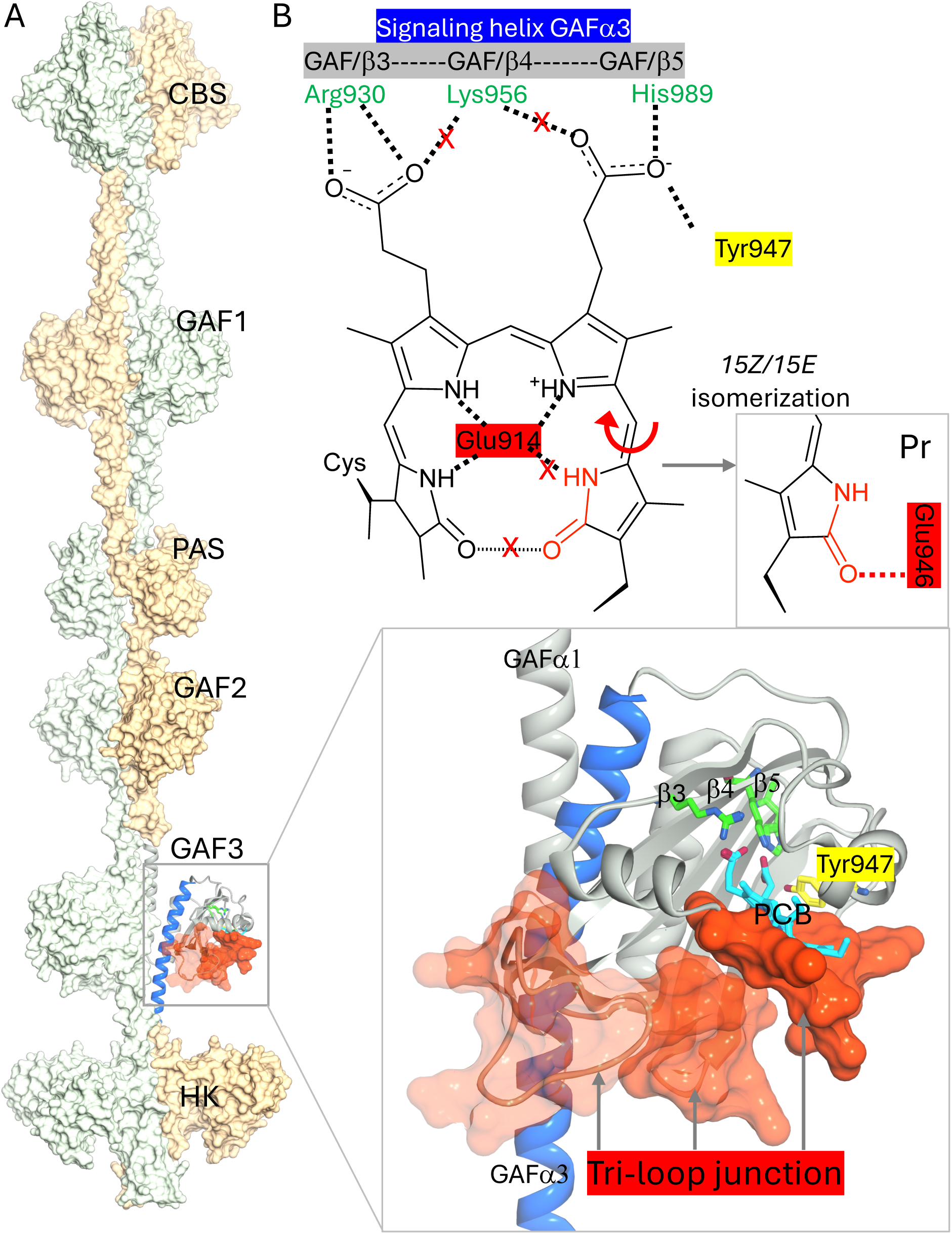
Proposed mechanisms for light signaling and spectral tuning in far-red CBCRs. **A)** AlphaFold2 model of multi-sensor Anacy2551 protein in a full-length dimeric scaffold where two protomers are shown in light yellow and light green surface, respectively. One bilin-binding GAF3 domain (i.e. 2551g3) is highlighted in a zoom-in view where the interactions between the GAF core residues (Arg930, Lys956 and His989 - green; Tyr947 - yellow) and propionates of the bilin chromophore (cyan) are highlighted. The tri-loop junction consisting of residues 908-916, 886-892 and 964-980 (red surface) offers a plausible pathway that transmits light-induced changes from the chromophore to the signaling helix GAFα3 at the dimer interface shared with the output histidine kinase (HK) domain. **B)** Key interactions responsible for inductive effects on spectral tuning. *15Z/15E* isomerization disrupts a short hydrogen bond between ring D nitrogen and Glu914 in the *15Z*-Pfr state. In one crystal form, the flipped ring D is stabilized by Glu946 giving rise to a *15E*-Pr state (gray box). In solution, 2551g3 proceeds to form the *15E*-Po state where interactions between the propionates and residues from the GAF core sheet are significantly weakened. See also Fig. S13.

This study also reveals a previously unrecognized distinction between light-absorbing states. The Pfr structure features a bilin chromophore tightly coupled with the GAF core through extensive interactions between positively charged residues coordinating both propionates. In contrast, these interactions are substantially weaker in the Po structure (Fig. 2, Fig. 5B). Accordingly, if chromophore affinity is inferred from the degree of protein-chromophore association observed by crystallography, the Pfr state may be viewed as a high-affinity ligand binding state whereas the Po state corresponds to a low-affinity state. By analogy to chemoreceptors where relatively small changes in ligand affinity are sufficient to alter signaling output, we propose that far-red CBCRs confers light signaling by reversibly switching the GAF domain between high-affinity Pfr state and low-affinity Po conformations. This model implies that the “flip-and-rotate” of the bilin chromophore, commonly observed in phytochromes and CBCRs, may represent a consequence, rather than a prerequisite of full photoconversion. Whether this affinity-based model widely applies to phytochromes that undergo the “flip-and-rotate” photocycle remains to be determined. Addressing this question will likely require experimental and/or computational methods capable of quantifying chromophore-protein affinity beyond static structural observations. For example, fluorescence-based methods might provide a means of probing the effective affinity of covalently bound chromophores in different signaling states. If validated, this affinity-based framework could provide a unifying mechanism across GAF-based signaling proteins, regardless of whether activation is initiated by light absorption or ligand binding.

## Materials and methods

### Protein purification and mutagenesis

The far-red CBCR construct 2551g3 consists of residues 839-1018 of Anacy2551, a multi-sensor histidine kinase protein from *Anabaena cylindrica* PCC 7122. The coding sequence of 2551g3 was inserted in the pET24b expression vector between the *NdeI* and *XhoI* restriction sites, resulting in a construct carrying a C-terminal hexa-histidine (6xHis) tag. 2551g3 was co-expressed in *E. coli* BL21(DE3) with a PCB biosynthesis plasmid encoding heme oxygenase 1 (HO1) and phycocyanobilin:ferredoxin oxidoreductase (PcyA). Cell growth, protein expression, and purification were performed as previously described (17).

### Crystallization and data collection of 2551g3 in the Po state

Crystals of 2551g3 in the Po state were grown under continuous white-light illumination at room temperature using the hanging drop vapor diffusion method. Purified protein at t 3 mg/mL was mixed with a 1:1 ratio with reservoir solution containing 25% (w/v) PEG 3350, 0.2 M ammonium sulfate, and 0.1 M bis-Tris pH 5.5 at room temperature. Prior to crystallization, protein samples were illuminated for 20 minutes with 750 nm ±20nm filtered light (Newport filter; ThorLabs OSL2 illuminator) to enrich the Po state. The identity of the Po state in the crystalline state was initially indicated by their characteristic blue coloration and subsequently verified by single-crystal absorption spectroscopy at room temperature (Fig. S3).

Single crystals of 2551g3-Po were harvested and cryo-protected using 25% glycerol as cryo-protectant. X-ray diffraction data were collected at 100 K at beamline 21-IDD of the Life Sciences Collaborative Access Team (LS-CAT) at the Advanced Photon Source (APS), Argonne National Laboratory. Diffraction images were indexed, integrated, and scaled using HKL2000 (54). The crystal structure of 2551g3-Po was determined in the I222 space group by molecular replacement (Phaser of CCP4) (55, 56) using the 2551g3 Pfr structure (PDB ID: 6UV8) as the search model.

Structure refinement was performed using Phenix (57). The final model was refined to 3.6 Å resolution with a final *R*-factor and free *R*-factor of 0.266 and 0.31, respectively.

### On-chip crystallization and Laue data collection in the dark-adapted state

Dark-adapted 2551g3 sample was crystallized on crystal-on-crystal devices using a batch method as previously described (58, 59). For each device, 5 μL of protein solution (10 mg/mL) was mixed with 5 μL of crystallization buffer containing 17% PEG 10,000, 0.1 M ammonium acetate, and Bis-Tris buffer (pH 5.5) in a 1:1 ratio at room temperature. The mixture was subsequently loaded onto a crystallization device and sealed between two quartz chips assembled with a 100-μm spacer. Crystals of varying sizes (typically 50-100 μm) appeared within 24 hours of setup. All crystallization devices were stored at room temperature in the dark until X-ray data collection.

Room-temperature serial Laue data were collected at beamline 14-IDB of BioCARS at APS using our automated high-throughput SerialX platform (23). To prevent unintended light activation, all crystallization devices were imaged exclusively under infrared illumination. Crystals were located. Ranked, and automatically introduced to the polychromatic X-ray beam for serial diffraction. All Laue diffraction images were indexed, refined, integrated and scaled using the Precognition/Epinorm^TM^ software package (Renz Research Inc). The structure was determined in the P4_2_ space group with two protein molecules in the asymmetric unit. The 2551g3-Pfr structure (PDB ID: 6UV8) was used as the search model for molecular replacement using Phenix/Phaser (57). The final structure was refined to 2.4 Å resolution using Phenix (57), yielding the final R-factor and R-free of 0.297 and 0.331, respectively. Model building and validation were performed in Coot, and structural figures were prepared using PyMOL (60) and Moorhen (https://moorhen.org).

### UV/visible spectroscopy in solution and single crystals

Solution spectra were recorded using Shimadzu UV-2600 spectrophotometer at room temperature. Absorption spectra of single crystals were measured by a microspectrophotometer equipped with a QE Pro spectrometer and a DH-2000-BAL deuterium-halogen light source (Ocean Insight). This custom-built instrument covers a broad spectral range of 200-1000 nm extending from mid-UV to near IR. Photoconversion of 2551g3 crystals was induced by far-red illumination provided by a LD785-SE400 laser diode (Thorlabs). Spectra were measured in time series using programmable timing control of both the illumination and spectrometer using a Raspberry Pi microcomputer via its versatile general-purpose input-output (GPIO) interface.

### Acid denaturation experiment and 15Z/15E composition analysis

Acid denaturation experiments were performed on both solution and crystal samples of 2551g3 under different illumination conditions to determine the *15Z/15E* composition. For solution samples, pre-illumination under green light (550 ± 30 nm) and red light (700 ± 30 nm) for 10-20 min to enrich Pfr and Po states, respectively. Acid denaturation solution (7 M urea, 150 mM NaCl, adjusted to pH 2.0 with concentrated HCl) was added to each sample followed by dark incubation on ice. The clarified denatured samples were used for UV–visible spectroscopy measurements.

For crystal samples, the 2551g3 crystals grown in dark on crystal-on-crystal chips were harvested, dissolved and denatured for spectroscopy measurements. Briefly, chips were observed under green filtered microscopic light before dissembled in a dark room for harvesting. Crystals transferred to a microcentrifuge tube were dissolved in low salt buffer (20mM Tris-HCl pH 8, 150mM NaCl) on ice followed by acid denaturation as described above. Low salt buffer was used for reference for UV/visible spectroscopic measurements on clarified denatured samples.

For *15Z/15E* composition analysis, least-squares fitting was performed using a six-parameter function defined as ***f***(x,y,t,u,v) = c0 + c1×x + c2×y + c3×t + c4×u - c5×v, which is implemented in Gnuplot (http://gnuplot.info). Denatured solution samples representing the *15Z* and *15E* states serve as basis spectra. Data points in a wavelength range of 250-900 nm were used to fit six parameters. Specifically, c0 and c1 are for baseline corrections of wavelength-independent and wavelength-dependent factors, respectively. c2 and c3 depict the *15Z*/*15E* composition while c4 and c5 are used for correcting discrepancy between reference spectra of solution and dissolved crystal samples. The overall goodness of fit is evaluated across the entire wavelength range (Fig. S3B).

### Construction of chimeric 2551g3HK and histidine kinase assays

The 604-residue chimeric construct 2551g3HK was cloned in the pET24b expression vector. Specifically, the photosensory domain (2551g3) consisting of Anacy2551 residues 828-1026 was fused to the catalytic histidine-kinase/receiver cassette (HKREC) containing PPHK residues 474-878 via an overlapping SacI site between the *NdeI–SacI* fragment of 2551g3 and *SacI–XhoI* fragment of PPHK. The resulting chimera carried a C-terminal hexa-histidine tag. For chromophore incorporation, 2551g3HK was co-expressed in *E. coli* BL21(DE3) with a PCB-producing plasmid, and protein purification was performed using the same protocols as described above.

PhosTag-based histidine kinase (HK) assays were performed according to established protocols developed for full-length PPHK (49). To evaluate the light-dependent kinase activities of 2551g3HK, filtered light at 550+/-20nm and 700+/-20nm was used to generate the Pfr and Po states, respectively. Protein samples were pre-illuminated, and continuous illumination was applied throughout the reaction to ensure the desired light-absorbing state. The isolated C-terminal receiver domain (cREC) from PPHK was used as a reporter protein as described previously.

### Vibrational spectroscopy

For spectroscopic experiments, dark-adapted 2551g3 in Tris buffer (pH 8.0) was concentrated to approximately 1 mM for RR and IR difference spectroscopy. The Pfr and Po states were generated by 3 - 5 minutes irradiation with a 580 and 780 nm laser diode, respectively. RR spectra were recorded using a Bruker Fourier-transform MultiRAM spectrometer equipped with a Ramanscope III module and a Nd-YAG CW laser at 1064 nm (line width, 1 cm^−1^; laser power at the sample, 680 mW) (Bruker, Karlsruhe, Germany). Measurements were performed at 90 K using a liquid-nitrogen-cooled cryostat (Linkam/Resultec). Spectra were typically accumulated for 1 h. Potential laser-induced damage was assessed by comparing spectra before and after a series of measurements, which revealed no detectable damage.

Infrared (IR) spectroscopic measurements were carried out at ambient temperature using a Bruker Tensor 27 FTIR spectrometer operated in transmission mode. Difference spectra were calculated as illuminated state minus initial state using a 1:1 subtraction ratio.

### Quantum mechanics/molecular mechanics (QM/MM) calculation

The crystal structure of 2551g3 (PDB ID: 6UV8) (17) served as the starting model for all computational studies, including vibrational spectrum simulations. To address potential steric clashes and optimize the structure, the model was first subjected to energy minimization using the ff14SB AMBER forcefield (61) as implemented in the AMBER program package (62, 63). For hybrid QM/MM optimization, the entire PCB chromophore and the side chain of the covalent anchor Cys943 were included in the QM region. This region was treated at the B3LYP/6-31G* level of theory with D3BJ dispersion correction (64–69). The remainder of the protein was described using molecular mechanics with the same force field used during the initial minimization. The QM/MM boundary was treated via hydrogen link atoms and interactions between the QM and MM regions was described using an electrostatic embedding scheme (70).

Vibrational frequency calculations were subsequently performed on the QM/MM-optimized structure with the same QM region and level of theory employed during geometry optimization. Harmonic vibrational frequencies were obtained by normal mode analysis (71). To account for systematic errors and facilitate comparison with experimental data, all frequencies were scaled by a factor of 0.96 (https://cccbdb.nist.gov/vibscalejustx.asp). To aid vibrational band assignments, the protons attached to the nitrogen atoms of pyrrole rings C and D were replaced with deuterons. Furthermore, anharmonic frequencies calculations were carried out using the same QM region and level of theory for better accuracy (72). The target vibrational modes included the C=O stretching vibrations of the bilin carbonyl groups. All QM/MM computations were performed using Gaussian16 (https://gaussian.com/gaussian16/).

## Acknowledgement

We thank the staff at the BioCARS and LS-CAT beamlines at Advanced Photon Source (APS) for their assistance in X-ray diffraction data collection. Use of the LS-CAT Sector 21 was supported by the Michigan Economic Development Corporation and the Michigan Technology Tri-Corridor under Grant 085P1000817. Use of BioCARS was supported by the National Institute of General Medical Sciences under Grant Number NIH P41 GM118217. Use of APS was supported by the U. S. Department of Energy (DOE) Office of Science, Office of Basic Energy Sciences, under Contract No. DE-AC02-06CH11357. This work was supported by National Institutes of Health grants R01EY024363 and R01EY035671 (to XY), and by Deutsche Forschungsgemeinschaft (DFG, German Research Foundation) grants 221545957 (CRC1072) and 390540038 (EXC 311 2008) (to PH).

## Author contributions

XY initiated and designed the project; LMB and SB carried out cloning, protein purification, and crystallization; AK and PH designed, performed and interpreted the RR and IR spectroscopic experiments; CW, AR and IS performed QM/MM calculations; LMB, SB and EN conducted mutagenesis; SB and XY collected monochromatic diffraction data; ZR and LMB collected Laue diffraction data; ZR processed the Laue data and carried out single-crystal spectroscopic experiments; XY determined, refined and analyzed crystal structures; XY, LMB, AK, PH, IS interpreted the data; XY, AK and PH wrote the paper with inputs from all authors.

The coordinates and X-ray diffraction data of dark-adapted and light-adapted structures have been deposited to the Protein Data Bank under the accession code of 10LA and 10KW, respectively.

