## Supplemental materials (1 table + 13 Figures) for "Crystal structures of a far-red photoreceptor in different light-absorbing states: insights into spectral tuning and light signaling"

### Table S1. Data collection and structure refinement statistics

| | 2551g3 (Dark-adapted state) | 2551g3 (Light-adapted state) |
|---|---|---|
| PDB ID | 10LA | 10KW |
| Diffraction data | | |
| Space group | $P4_2$ | I222 |
| a, b, c (Å) \| α, β, γ (°) | 108.8, 108.8, 71.1 \| 90, 90, 90 | 151.6, 353.6, 67.1 \| 90, 90, 90 |
| Data collection Method | In situ serial Laue diffraction | Monochromatic oscillation |
| Data collection temperature | Room temperature (~298 K) | 100 K |
| Beamline | 14-IDB, BioCARS, APS | 21-IDD, LS-CAT, APS |
| # of images | 3936 indexed Laue images | 180 1-deg oscillation images |
| Resolution (Å) | 50.00 – 2.4 (2.5-2.4) | 50-3.6 (4.2-3.6) |
| ***R***merge | 0.045 (0.041) | 0.084 (0.775) |
| Completeness (%) | 92.5 (85) | 99.7 (99.7) |
| Redundancy | 420 (450) | 6.2 (5.5) |
| I/σ(I) | 1.1 (0.7) | 11 (1.1) |
| Asymmetric unit | | |
| Protein | Chains A, B (180 aa/chain) | Chains A, B, C, D, E (180 aa/chain) |
| Waters | 23 | |
| Ligands | 2 CYC | 5 CYC |
| Refinement | | |
| Resolution (Å) | 20-2.4 (2.45-2.4) | 46-3.6 (3.79-3.6) |
| R-factor | 0.297 (0.390) | 0.246 (0.437) |
| free R-factor | 0.331 (0.429) | 0.297 (0.447) |
| RMSD bond length (Å) | 0.009 | 0.003 |
| RMSD bond angle (°) | 1.121 | 1.113 |
| Average B ($Å^2$) | 9.59 | 134.7 |
| Ramachandran | | |
| Favored (%) | 95.6 | 95 |
| Allowed (%) | 4.4 | 4.8 |
| Disallowed (%) | 0 | 0.2 |

SI Fig. S1

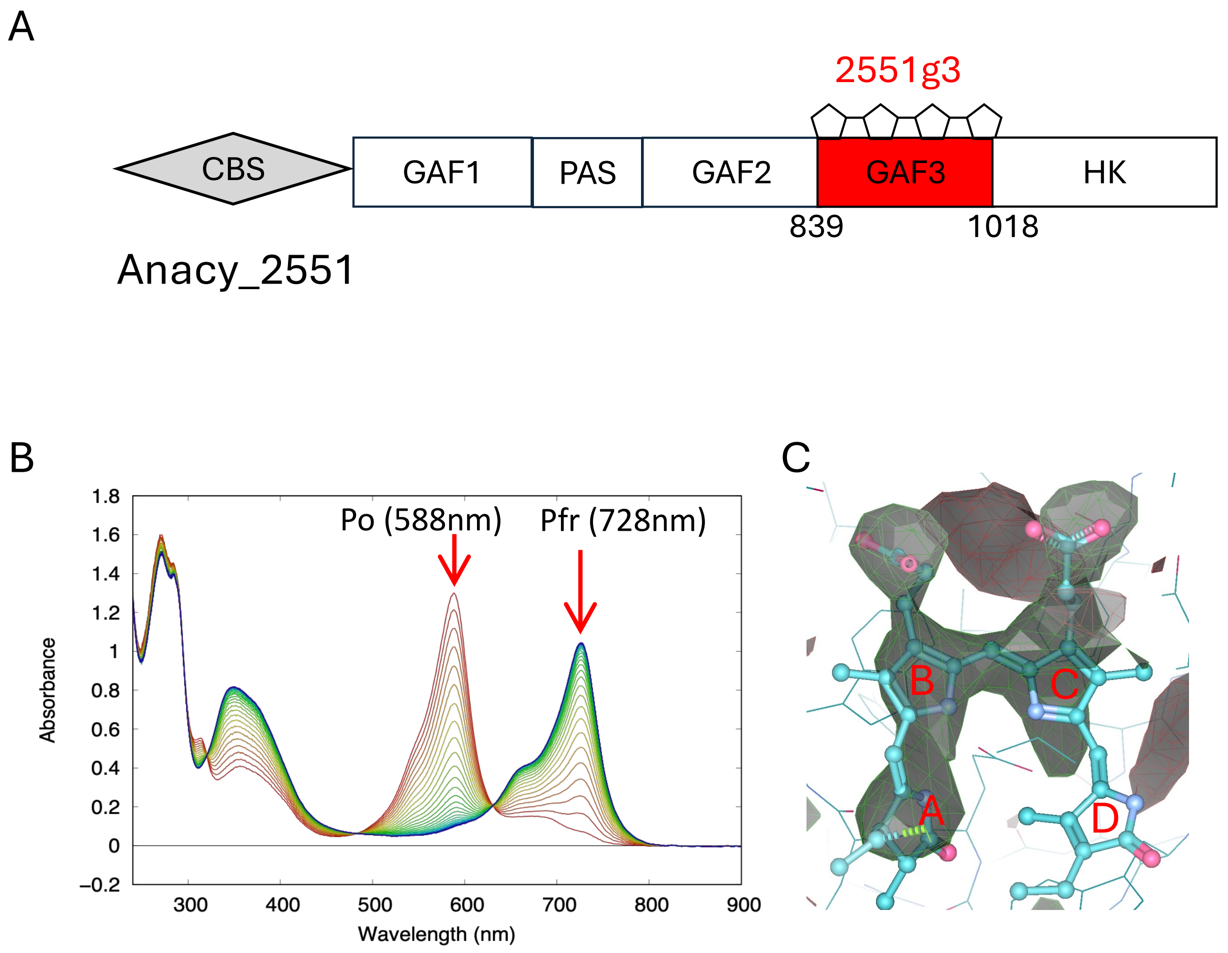

**Figure S1. Anacy_2551g3 is a representative far-red cyanobacteriochrome (CBCR). A)** Domain structure of full-length multi-domain histidine kinase Anacy_2551. Anacy_2551g3 (2551g3 in short) used in this study is the isolated GAF3 domain consisting residues 839-1018. **B)** Time series of 2551g3 photoconversion in solution between the Pfr (peak at 728 nm) and Po (peak at 588 nm). **C)** Maps obtained from cryo-crystallography show highly disordered electron densities for ring D of the bilin chromophore, due to inadvertent often inevitable light exposure during crystal transfer and harvesting.

SI Fig. S2

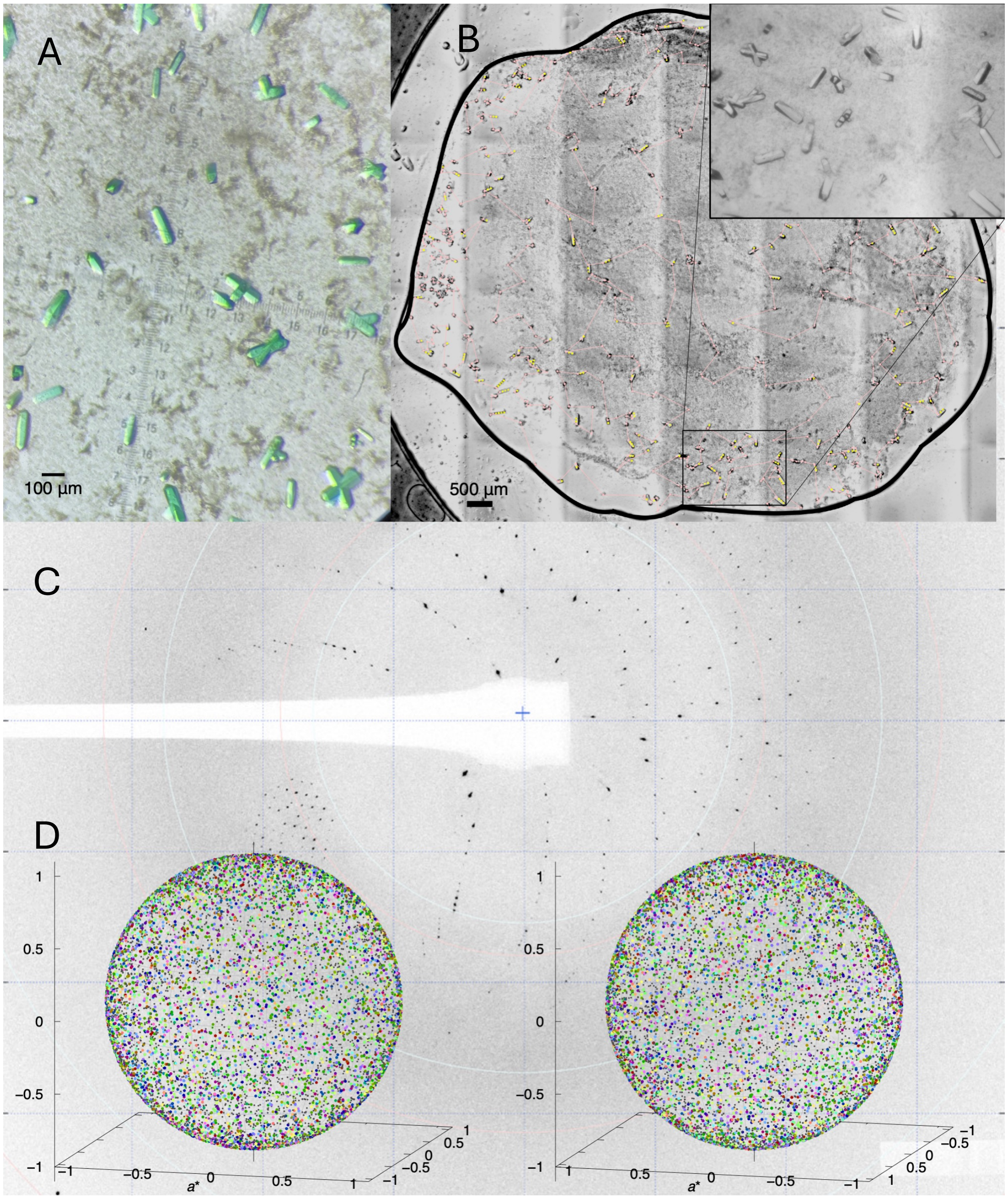

**Figure S2. . Serial Laue data collection using *in situ* serial crystallography at room temperature. A**) Green 2551g3 crystals under microscope. **B)** A tiled image of the entire crystallization drop viewed under infrared light. Crystals used for X-ray shots are marked by small yellow circles. The inset shows a zoom-in view of crystal grown on the device. **C)** A typical Laue diffraction pattern from *in situ* diffraction. **D)** Distribution of crystal orientation from nearly 4000 crystals. Crystal orientations represented by colored dots on two hemispheres are derived from Laue image indexing. Crystallization devices are distinguished by color.

SI Fig. S3

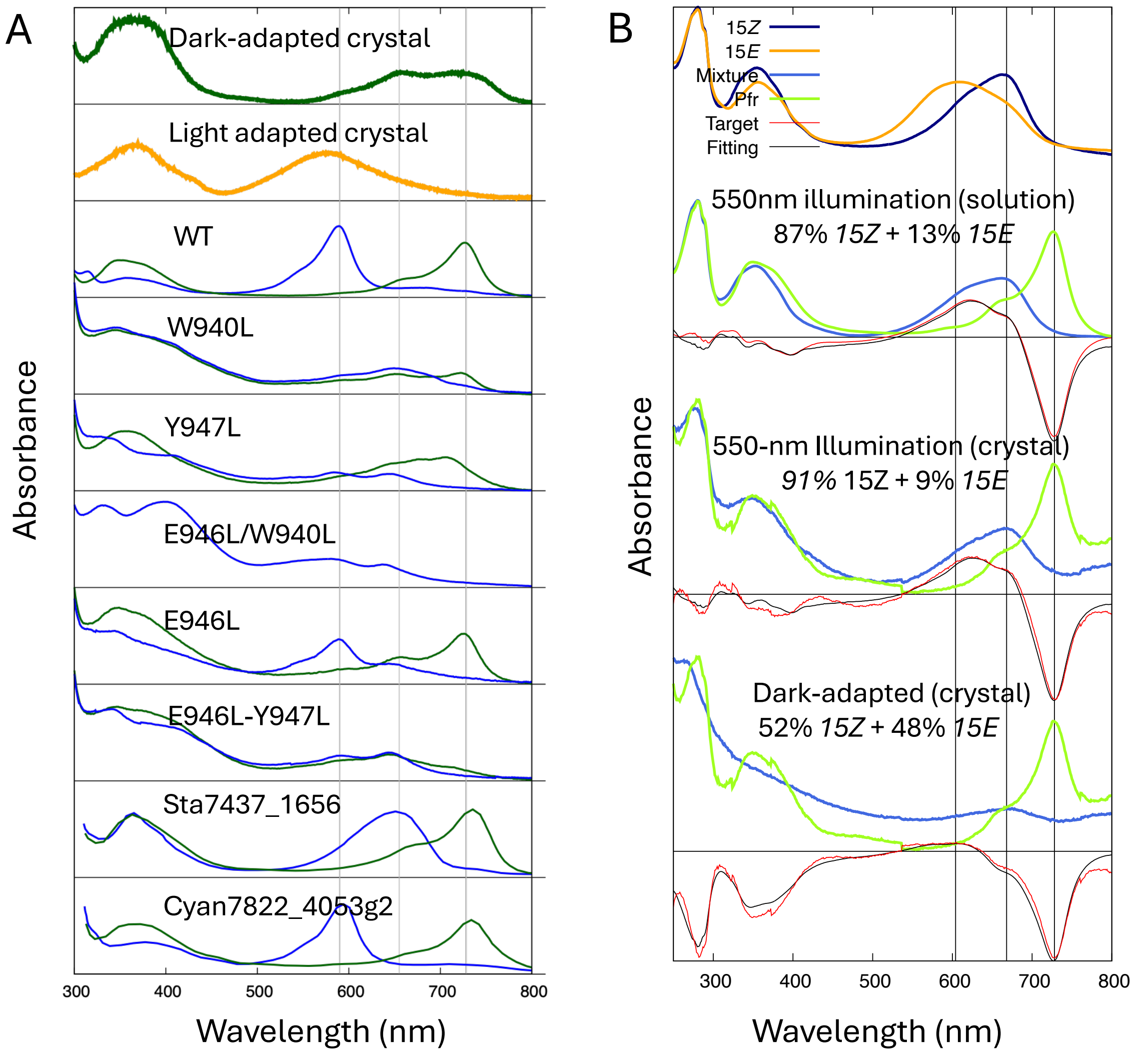

**Figure S3. Spectral analysis of 2551g3. A)** Absorption spectra of 2551g3 single crystals and mutants in solution. Spectra of wild type (WT) 2551g3 and two homologous far-red CBCRs (Sta7437_1656 and Cyan7822_4053g2) are also for references (Rockwell et al, 2015). Thin gray lines mark peak wavelengths at 588, 655 and 728 nm, corresponding to the Po, Pr and Pfr states, respectively. **B)** Compositional analysis. With two basis spectra (top panel) representing the denatured *15Z* (dark blue) and *15E*-Po (orange) states, spectra of denatured solution and crystal samples (blue) are subjected to baseline correction and least-squares fitting for determining the *15Z/15E* composition (see Methods). Reference spectra used for calculating target difference spectra are shown in green. The goodness of fit is supported by overall agreement between fitted (black) and target (red) difference spectra. Thin lines mark wavelengths at 604, 668 and 728 nm, corresponding to the denatured *15E*, denatured *15Z* and Pfr states, respectively.

### SI Fig. S4

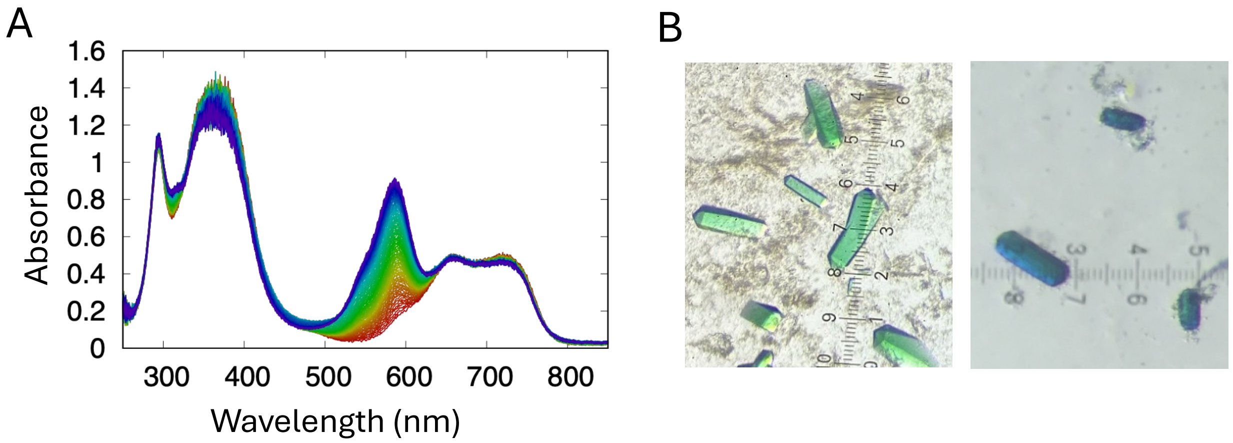

**Figure S4. Photoactivity in single crystals. A)** Time series of absorption spectra (from red to blue) from a single crystal of 2551g3 are recorded under continuous 785-nm illumination. **B)** Left: 2551g3 dark-adapted crystals visualized under white microscopic light. Right: Blue-colored crystals are observed after filtered far-red (785-nm) light illumination for 5-10 minutes.

SI Fig. S5

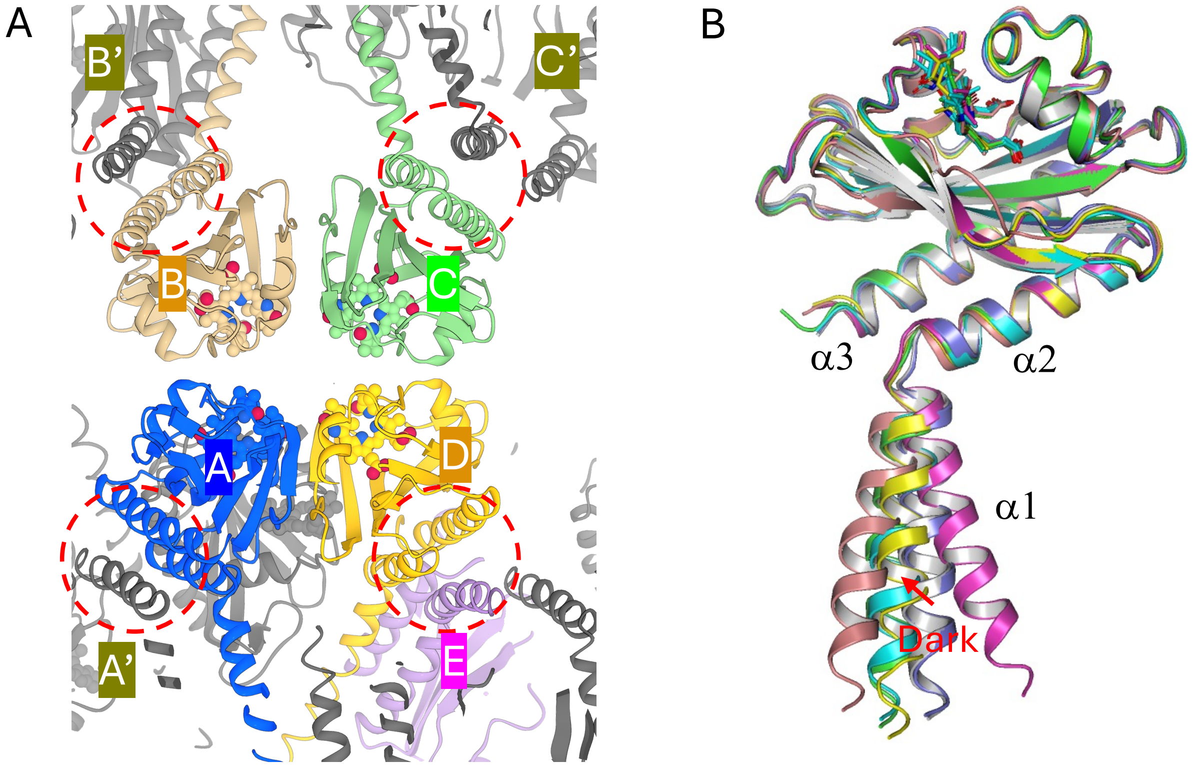

**Figure S5. Crystal structure and molecular packing of light-adapted 2551g3 in space group I222. A)** Five molecules in asymmetric unit (ASU) form a network of dimers via intermolecular 3-helix bundles (marked in red circles). Chains A-C dimerize via crystallographic 2-fold symmetry while chains D-E are related by non-crystallographic 2-fold symmetry. Molecules within ASU are shown in color while crystallographic symmetry mates are shown in dark gray. **B)** Structural alignment according to the GAF core reveals large variation in the disposition of $\alpha 1$ helices among five protomers. The corresponding $\alpha 1$ helices of the dark-adapted structure are also shown (gray and blue) for comparison.

### SI Fig. S6

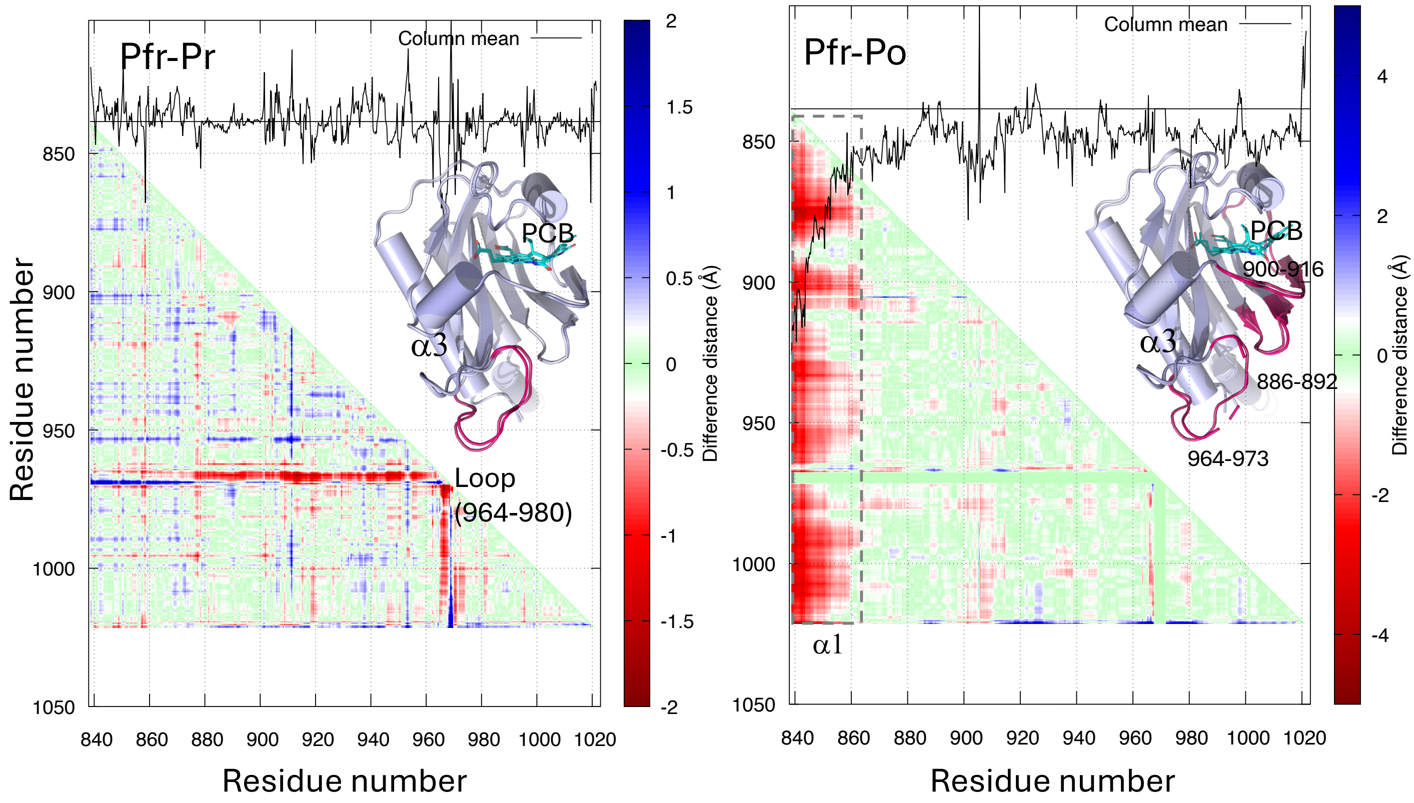

**Figure S6. Protein structural differences between the Pfr, Pr and Po states. A)** Difference distance matrix between the main chain atoms of *15Z*-Pfr and *15E*-Pr subunits. Amplitude of change is coded according to color bar (right). The most notable differences reside within the large β4-β5 loop insertion spanning residues 964-973 (red color in ribbon diagram). **B)** Difference distance matrix between the main chain atoms of *15Z*-Pfr and representative *15E*-Po structure. The overall amplitude of change is larger compared to the left panel. In addition to GAF-α1 (marked by dashed line), three segments (residues 886-892, 900-916, 964-973, red color in ribbon diagram) show prnounced changes. They correspond to three loops that constitute the tri-loop junction. Characteristic to far-red CBCRs, the tri-loop junction bridges the chromophore (cyan) to the GAF-α3 helix (Bandara et al, 2021).

SI Fig. S7

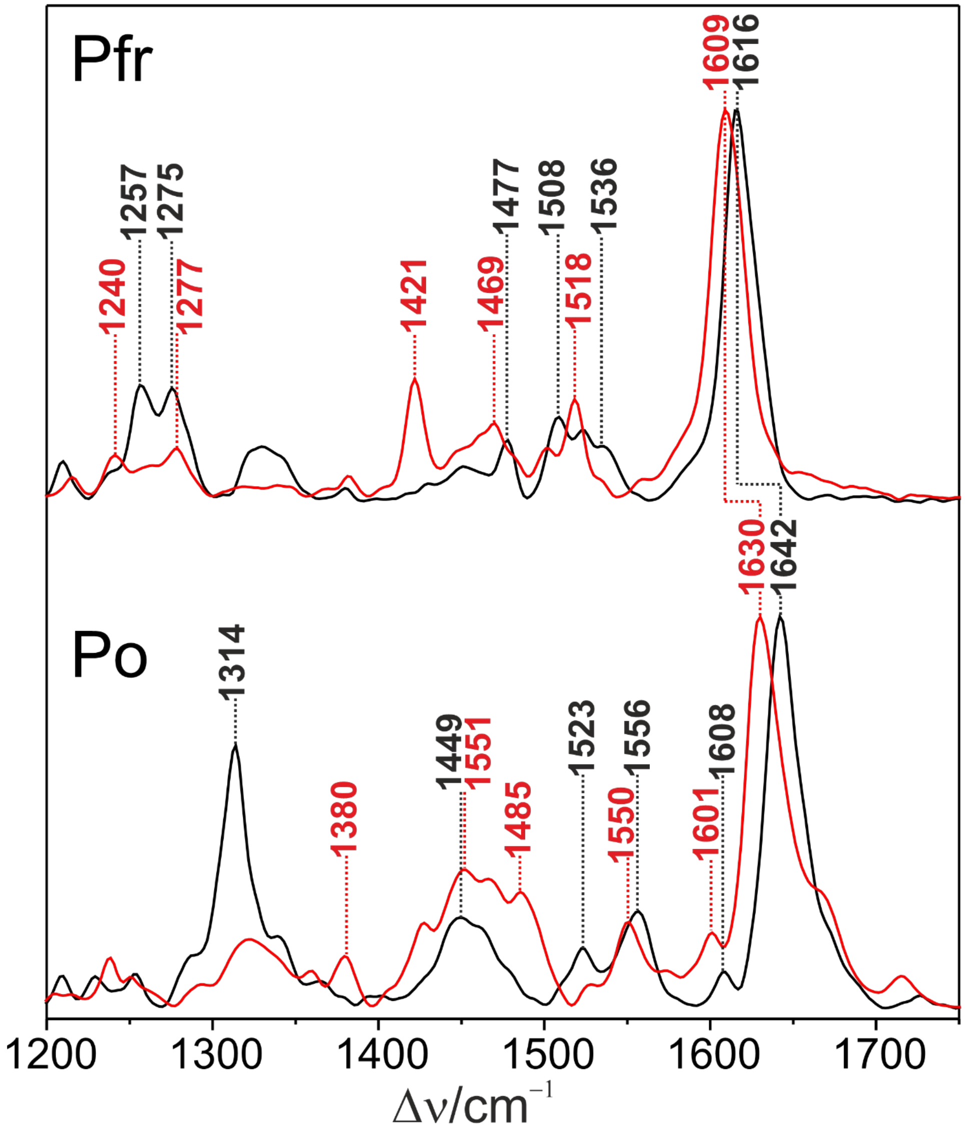

**Figure S7. Resonance Raman spectra of 2551g3 in the Pfr and Po states** obtained in $H_2O$ (black) and $D_2O$ (red) at pH (pD) = 8.0. The spectra were measured with 1064 nm excitation at 90 K.

SI Fig. S8

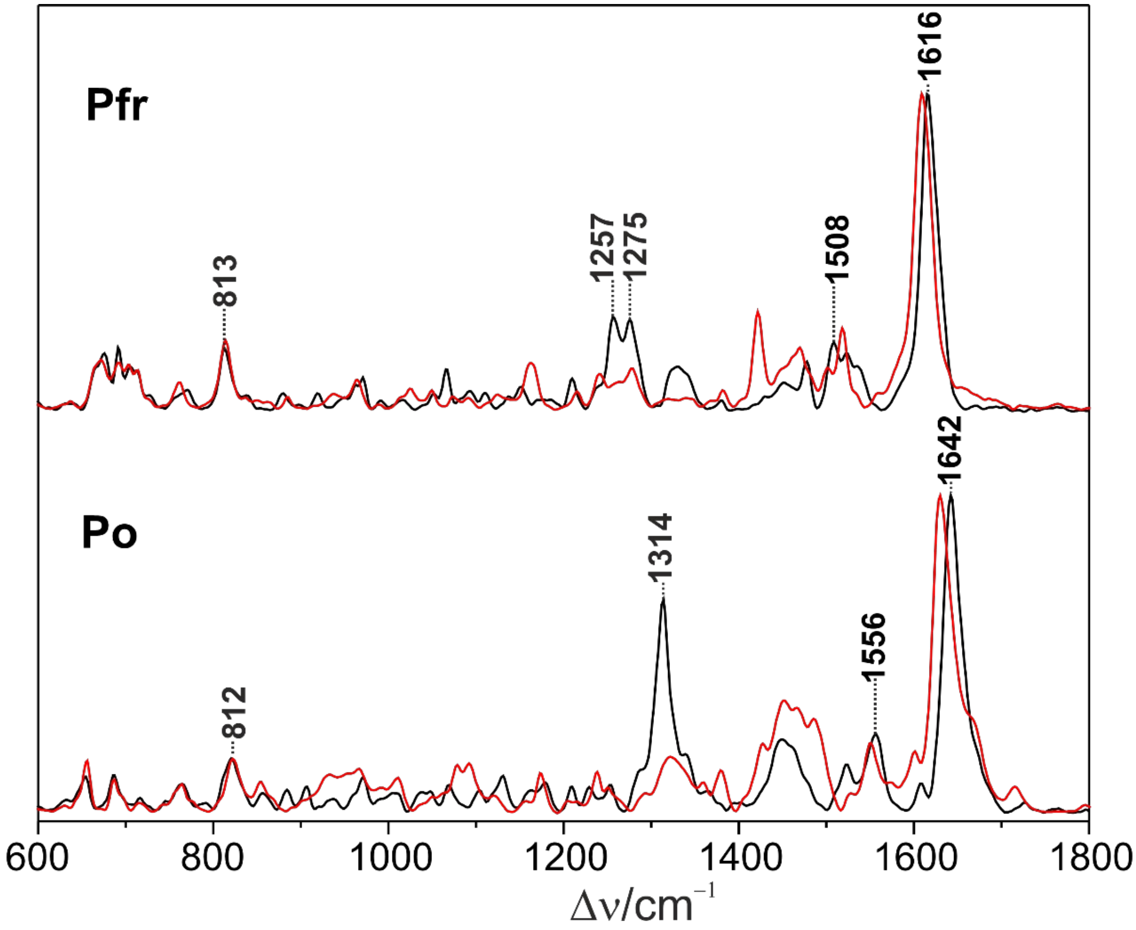

**Figure S8. Resonance Raman spectra of 2551g3 in the Pfr and Po states** obtained in $H_2O$ (black) and $D_2O$ (red) at pH (pD) = 8.0. The spectra were measured with 1064 nm excitation at 90 K.

SI Fig. S9

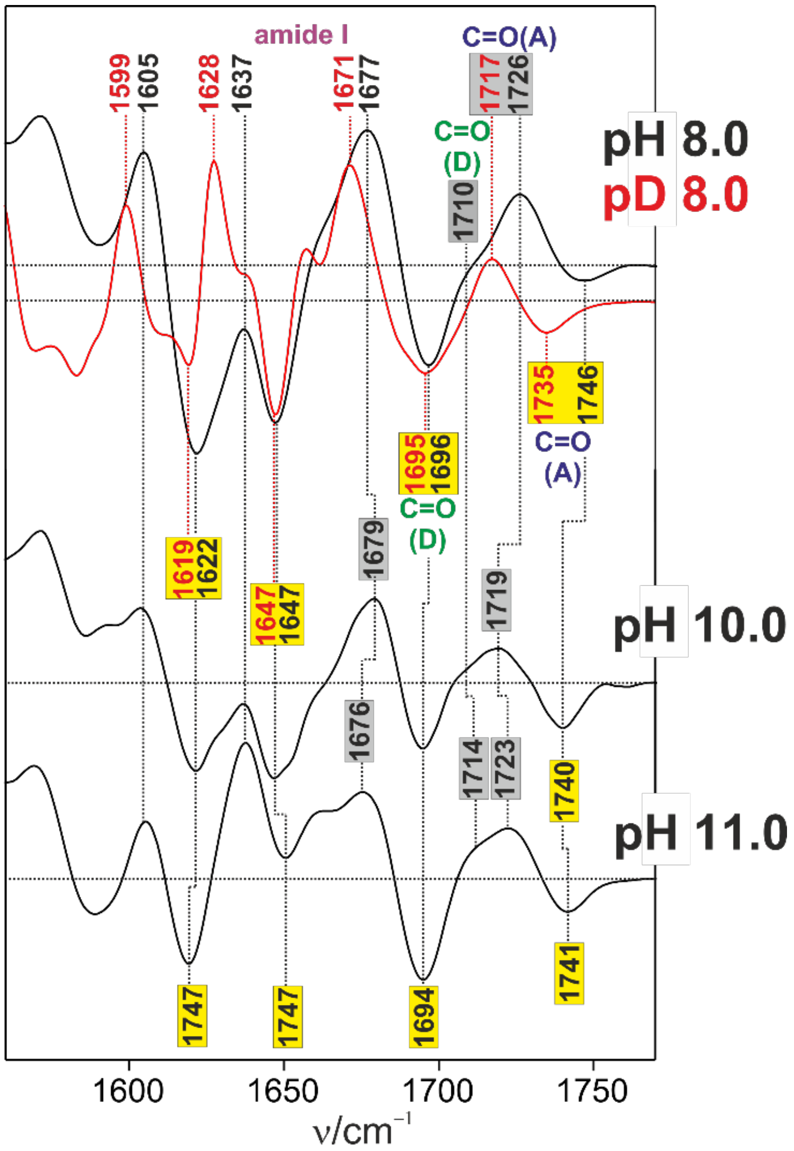

**Figure S9. "Po minus Pfr" IR difference spectra measured at different pH.** Black and red traces refer to $H_2O$ and $D_2O$, respectively. Positive and negative signals in the difference spectra originate from Po and Pfr, respectively.

Fig. S10

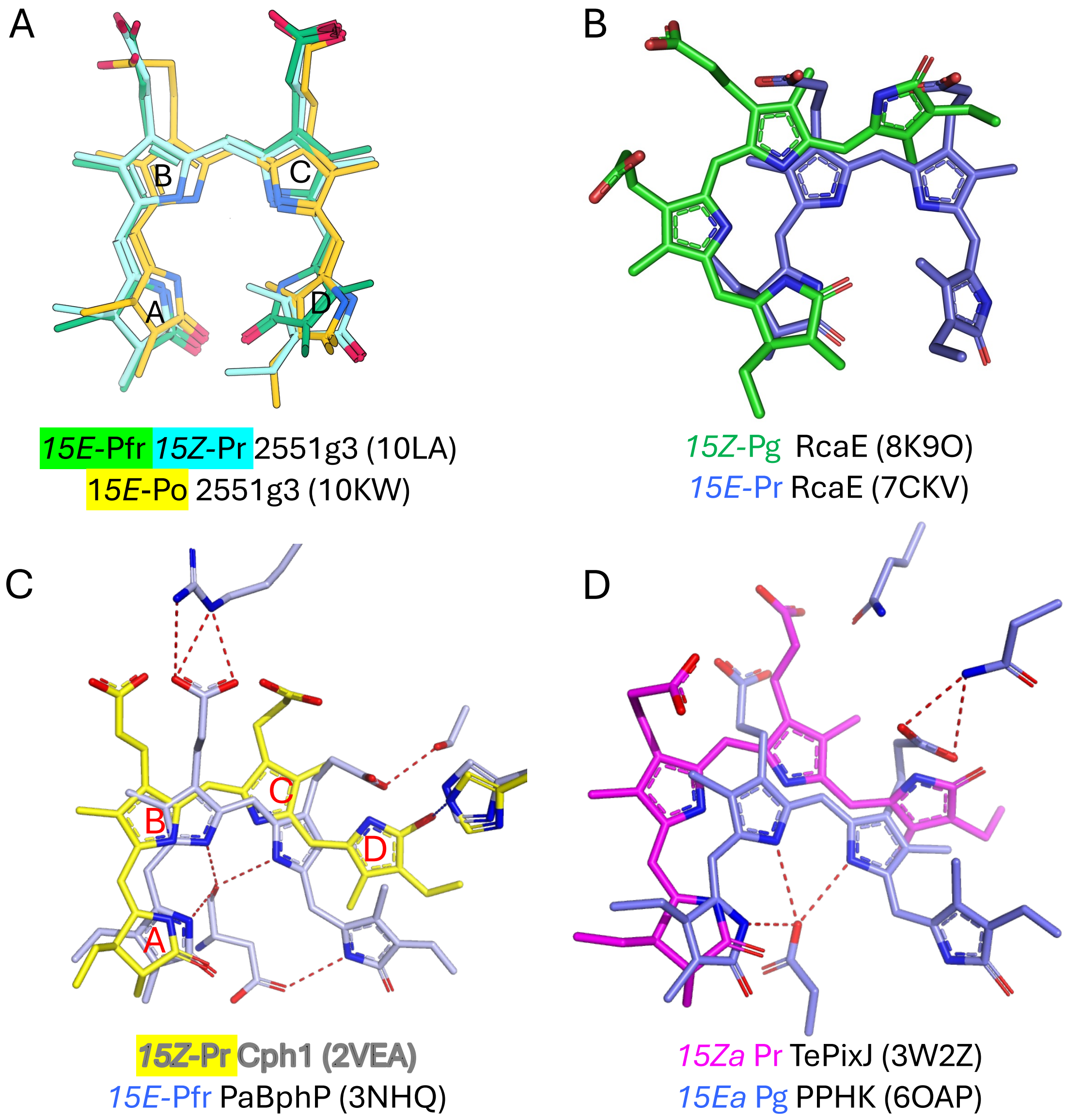

**Figure S10. Photoconversion of 2551g3 does not involve the flip-and-rotate motion of the bilin chromophore. A)** Superposition of the bilin chromophores based on protein structural alignment via SSM (secondary structure match). Ring D flip involves little chromophore rotation relative to the GAF core. **B)** The Pr and Pg structures of RcaE shows a large rotational motion of the PCB chromophore relative to the GAF core. **C,D)** Representative bacteriophytochromes and CBCRs show typical flip-and rotate" motions associated with photoconversion. PDB accession codes used for structural comparisons are shown in parentheses.

SI Fig. S11

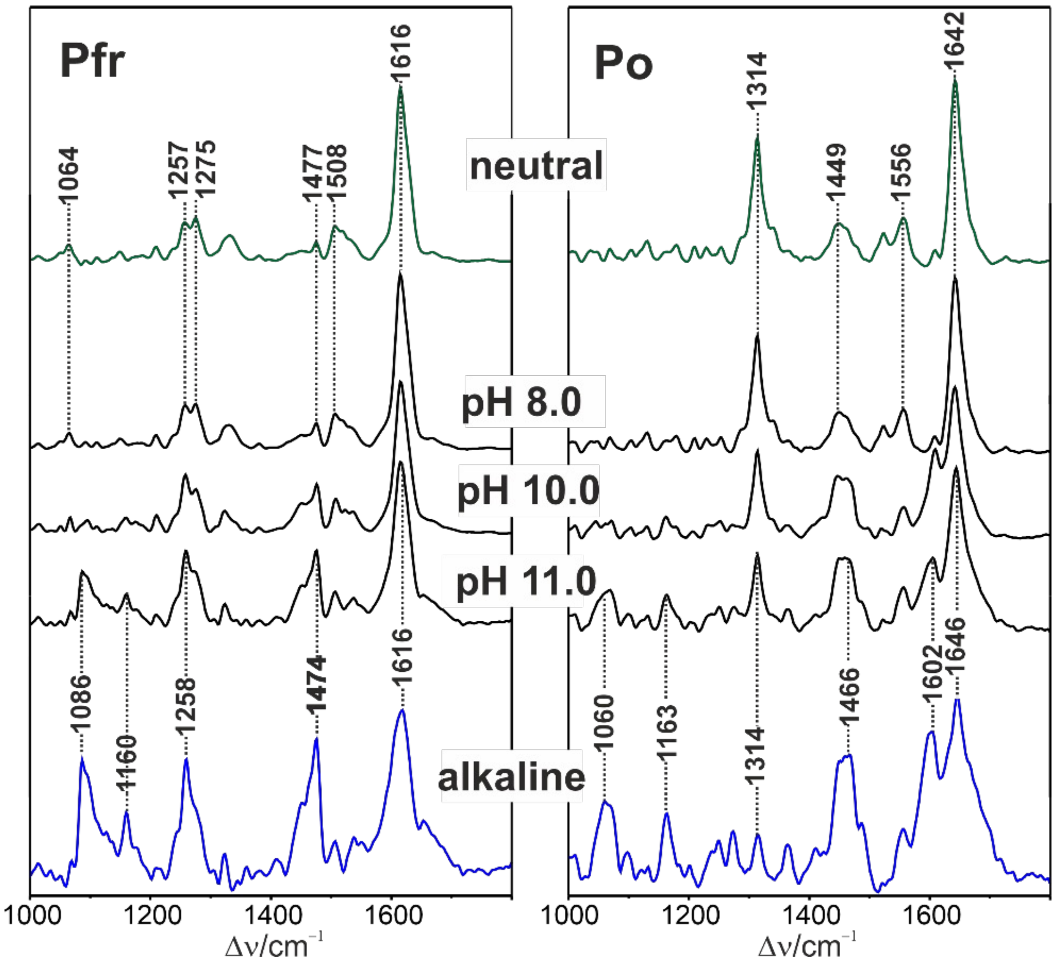

**Figure S11. RR spectra measured at different pH (black) of Pfr (left) and Po (right).** The green (top) and blue (bottom) traces are difference spectra obtained by subtracting the spectra at pH 11 from those at pH 8 (top) and vice versa (bottom), leading to the so-called neutral and alkaline forms, respectively. The subtraction factors for the alkaline forms are 0.45 and 0.50 for Pfr and Po, respectively.

SI Fig. S12

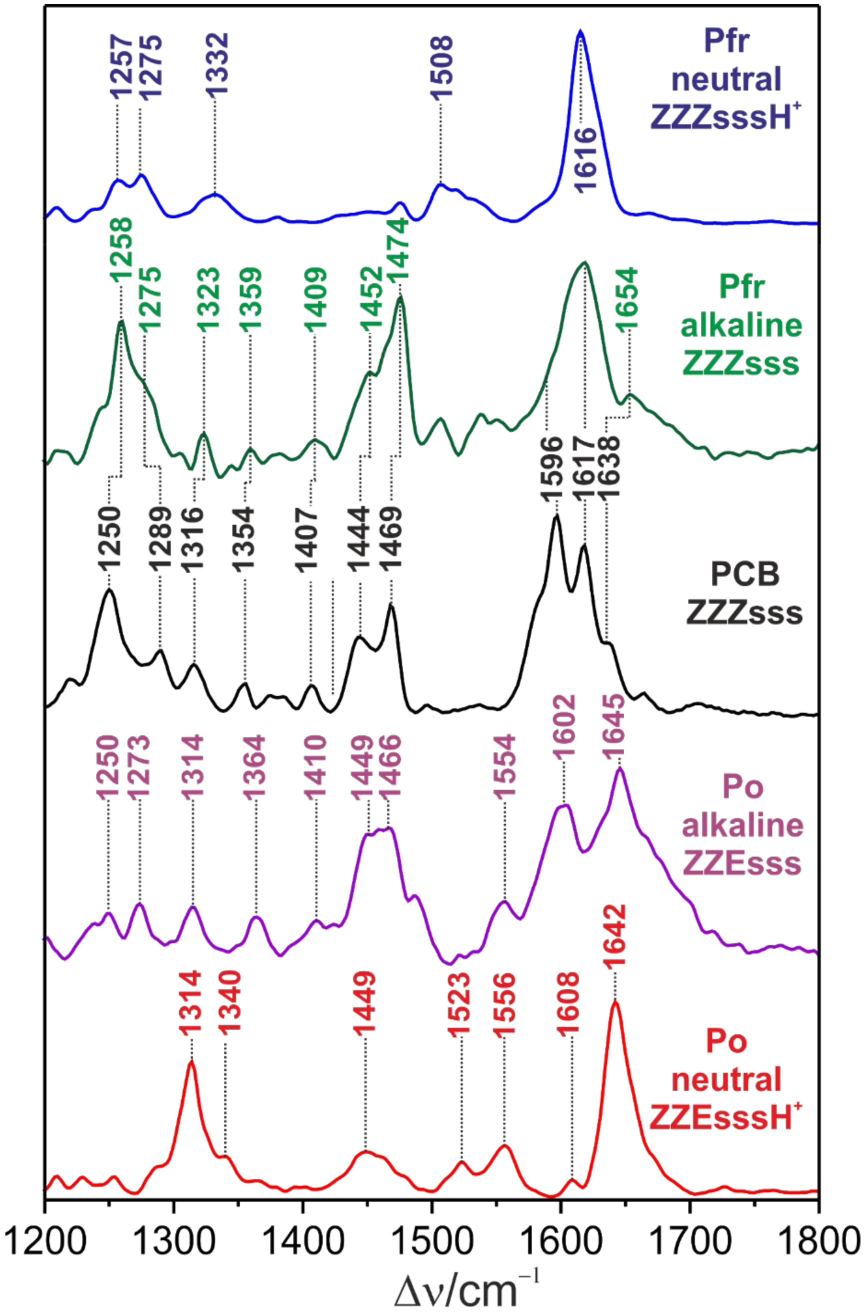

**Figure S12. Pure RR spectra of the neutral forms of Pfr (blue) and Po (red).** The alkaline forms of Pfr (green) and Po (magenta) were obtained by mutual subtraction of the corresponding spectra measured at pH 8 and pH 11 (see Fig. S8), and the spectra of solid PCB in the *ZZZsss* configuration (black).

SI Fig. S13

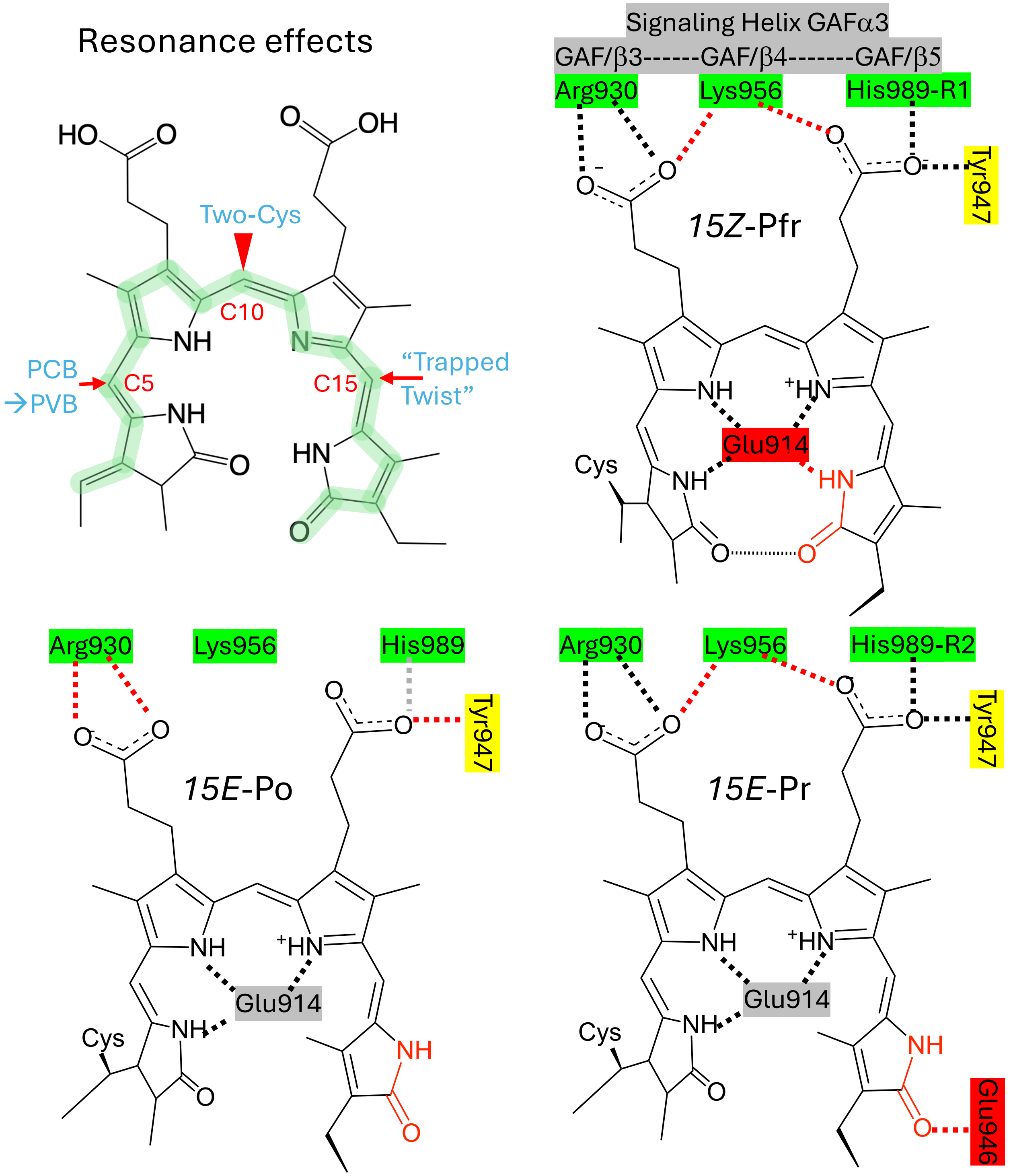

**Figure S13. Mechanisms of spectral tuning.** Top-left: Three examples of resonance effects that directly acts on the bilin backbone (red arrows) in bilin-based photoreceptors. Protein-chromophore interactions observed in this work highlight different electrostatic or inductive effects in three light-absorbing states. His989-R1 and His989-R2 mark different rotamers in the Pfr and Pr states, respectively. Interactions distinct between states are highlighted in red. While *15Z*-Pfr and *15E*-Pr exhibit close association between the GAF core and propionates, the *15E*-Po state show weakened interactions between the chromophore and GAF central β strands.
